# A 6.5kb intergenic structural variation enhances P450-mediated resistance to pyrethroids in malaria vectors lowering bed net efficacy

**DOI:** 10.1101/2020.05.05.078600

**Authors:** Leon M.J. Mugenzi, Benjamin D. Menze, Magellan Tchouakui, Murielle J. Wondji, Helen Irving, Micareme Tchoupo, Jack Hearn, Gareth D. Weedall, Jacob M. Riveron, Fidelis Cho-Ngwa, Charles S. Wondji

## Abstract

Elucidating the complex evolutionary armory that mosquitoes deploy against insecticides is crucial to maintain the effectiveness of insecticide-based interventions. Here, we deciphered the role of a 6.5kb structural variation (SV) in driving cytochrome P450-mediated pyrethroid resistance in the malaria vector, *Anopheles funestus*. Whole genome pooled sequencing detected an intergenic 6.5kb SV between duplicated CYP6P9a/b P450s in pyrethroid resistant mosquitoes through a translocation event. Promoter analysis revealed a 17.5-fold higher activity (P<0.0001) for the SV-carrying fragment than the SV-free one. qRT-PCR expression profiling of *CYP6P9a/b* for each SV genotype supported its role as an enhancer since SV+/SV+ homozygote mosquitoes had significantly greater expression for both genes than heterozygotes SV+/SV- (1.7-2-fold) and homozygotes SV-/SV- (4-5-fold). Designing a PCR assay revealed a strong association between this SV and pyrethroid resistance (SV+/SV+ vs SV-/SV-; OR=2079.4, *P*=<0.001). The 6.5kb SV is present at high frequency in southern Africa (80-100%) but absent in East/Central/West Africa. Experimental hut trials revealed that homozygote SV mosquitoes had significantly greater chance to survive exposure to pyrethroid-treated Nets (OR 27.7; P < 0.0001) and to blood feed than susceptible. Furthermore, triple homozygote resistant (SV+/CYP6P9a_R/CYP6P9b_R) exhibit a higher resistance level leading to a far superior ability to survive exposure to nets than triple susceptible mosquitoes, revealing a strong additive effect. This study highlights the important role of structural variations in the development of insecticide resistance in malaria vectors and their detrimental impact on the effectiveness of pyrethroid-based nets.

## Introduction

Malaria control programs rely heavily on insecticide-based vector control interventions including the mass distribution of long lasting insecticidal nets (LLINs) impregnated with pyrethroids [1]. Unfortunately, the increased use of LLINs over the years has contributed, among other factors, to the selection of pyrethroid resistant mosquitoes [2]. The widespread of resistance to insecticides in major malaria vectors including *An. gambiae* and *An. funestus* is probably one of the main factors behind the recent increase in malaria cases across the world, from 214 million in 2015 to 219 million in 2017 [3] or the stagnation of such control effort [4]. Such growing resistance reports calls for urgent action to implement suitable resistance management strategies to reduce the impact on the effectiveness of current and future insecticide-based tools as highlighted by the WHO global plan for insecticide resistance management [5]. Elucidating the molecular and genetic basis of insecticide resistance is a key step to understanding factors driving resistance and to design diagnostic tools to better detect and track the spread of such resistance in the field as well as better assess its impact on the effectiveness of control tools [5].

The two major insecticide resistance mechanisms are target-site resistance and metabolic resistance. Target-site resistance mechanisms including the knockdown resistance (*kdr*) in the sodium channel gene are well characterized in most mosquito species [6]. However, for other species such as *An. funestus, kdr* has been shown to be absent with resistance mainly conferred by metabolic resistance [7]. Three classes of enzymes are mainly conferring metabolic resistance notably cytochrome P450 monooxygenases (P450s), glutathione S-transferases and esterases [8]. However, the molecular drivers of metabolic resistance have been more difficult to decipher due to the greater complexity of this mechanism with several genes involved and various molecular processes potentially implicated. Nevertheless, progress has been made recently in elucidating the molecular basis of metabolic resistance in malaria vectors. For example, amino acid changes associated with increased metabolic activity of detoxification genes have been detected notably in the *GSTe2* gene with L119F detected in *An. funestus* [9] and I114T in *An. gambiae* [10]. The L119F-GSTe2 was confirmed to confer cross resistance to pyrethroids and DDT [9]. An allelic variation of detoxification genes has also been shown to confer insecticide resistance as described in the case of the P450 genes *CYP6P9a* and *CYP6P9b* [11] and for *GSTe2* in the dengue vector *Aedes aegypti* [12]. Recently, causative mutations located in the cis-regulatory region of key P450 genes were shown to drive the over-expression of the P450s *CYP6P9a* [13] and *CYP6P9b* [14] in *An. funestus*. Moreover, evidence that copy number variation of detoxification genes was also playing a role in the observed insecticide resistance in mosquitoes was shown in the *An. gambiae* and *An. coluzzii* [15]. Additionally, key transcription binding factors notably Maf-S/CncC [13, 16] have been associated with insecticide resistance in major malaria vectors. In the case of *An. funestus*, a major structural variation (SV) in the shape of a 6.5kb insertion was reported in pyrethroid resistance mosquitoes in the intergenic regions of two P450s (*CYP6P9a* and *CYP6P9b*) [13] raising the prospect that such structural variation could also be an key factor driving metabolic resistance in mosquitoes. Establishing the potential role of such structural variations could help elucidate the molecular basis of resistance and also design robust diagnostic tools to detect and track resistance in the field for current and future insecticides.

Therefore, to elucidate the potential role of structural variation in pyrethroid resistance, we assess the contribution of the 6.5kb insertion at the vicinity of key P450 resistance genes in resistant *An. funestus* mosquitoes. We demonstrate using functional assays and genotype/phenotype studies that the 6.5kb is playing a key role in the over-expression of P450 resistance genes. We designed a PCR-based diagnostic that allows to track this SV-based resistance and use it to show that this 6.5kb acts as an enhancer significantly contributing to increase pyrethroid resistance and to exacerbate the loss of efficacy of long-lasting insecticidal nets against malaria vectors.

## RESULTS

### Identifying the presence or absence of the insertion in samples from different geographical locations in Africa

By aligning the intergenic region between *CYP6P9a* and *CYP6P9b* for FUMOZ and FANG, the insertion point was identified at 851 bp from the stop codon of *CYP6P9b* and 840 bp from the start codon of *CYP6P9a*. To determine the geographical extent of the 6.5k SV across African populations of *An. funestus*, pooled-template whole genome sequencing alignments to the 120kb *rp1* BAC from several locations across the range of *An. funestus* were inspected. This *rp1* BAC spans the cytochrome P450 cluster where both *CYP6P9a* and *CYP6P9b* are located [17]. Fig 1 shows alignments for FUMOZ-R and FANG, which show both anomalous features. S1A and S1B Fig shows the putative left and right ends of the insertion, defined by the presence of “clipped” reads, where the alignment software aligns a part of the read to the reference and “clips” off part of the read that does not align. The FUMOZ-R alignment contains reads that are left-clipped (the leftmost part of the read, as aligned to the reference, is clipped irrespective of the read’s orientation: so the 5’ end of a read aligned to the positive strand and the 3’ end of a read aligned to the negative strand) between BAC sequence positions 37409 and 37410 (37410 being the leftmost base included in the insertion). It also contains reads that are right clipped between positions 43954 and 43955. This defines a region of 6545 bp. The presence of only left-clipped reads on the left of the region and right-clipped reads on the right of the region indicates two things in FUMOZ-R: (i) that the “insertion” form of the indel is fixed in the FUMOZ-R sample (there is no evidence for presence of the “deletion” form), and (ii) that the inserted sequence is homologous to part of a larger sequence found elsewhere in the genome (indicated by the “clipped” parts of the reads). In the susceptible FANG the situation is more complicated. At the left end of the insertion there are reads left-clipped between positions 37409 and 37410 (as for FUMOZ-R) but also some reads right-clipped slightly further left, between positions 37404 and 37405. At the right end of the insertion there are reads right-clipped between positions 43954 and 43955 (as for FUMOZ-R) but also some reads left-clipped between the same positions. In addition, further to the right there are some reads right-clipped between positions 44053 and 44054 and some left-clipped between positions 44070 and 44071. Detailed inspection of the clipped reads showed that the reads right-clipped at 37404/37405 and left-clipped at 43954/43955 indicate the “deletion” form of the indel, as the clipped parts of the reads from the left and right end of the insertion overlap each other (but also contain a short length of DNA that does not match FUMOZ-R). The clipping at 44053/44054 and 44070/44071 is due to a region of 35 bp in FANG (TAA TAC CGG GAG ATA CAT GGA GCT CGT GTA AAA GA) that does not align with the FUMOZ reference (ATA TGT CGG AGG TTT AT) at the same location. Overall, FANG shows evidence of the “deletion” form of the indel in addition to the presence of the large homologous sequence elsewhere in the genome (Fig 1A). This makes simple inspection of the alignment misleading, as rather than a loss of coverage across the 6.5 kb indel, coverage is seen due to reads originating from sequence elsewhere in the genome. This is illustrated in S1A Fig. Overall, FUMOZ-R exhibits an increased coverage depth of reads between *CYP6P9a* and *CYP6P9b* in with a loss of polymorphism compared to the susceptible FANG strain (Fig 1B). The same contrast is observed between the 2002 sample from Mozambique before the scale up of LLIN (2002) with a low coverage of read depth contrary to the 2016 sample showing a greater coverage depth at the intergenic region corresponding to the insertion of the 6.5kb fragment.

**Figure 1:**
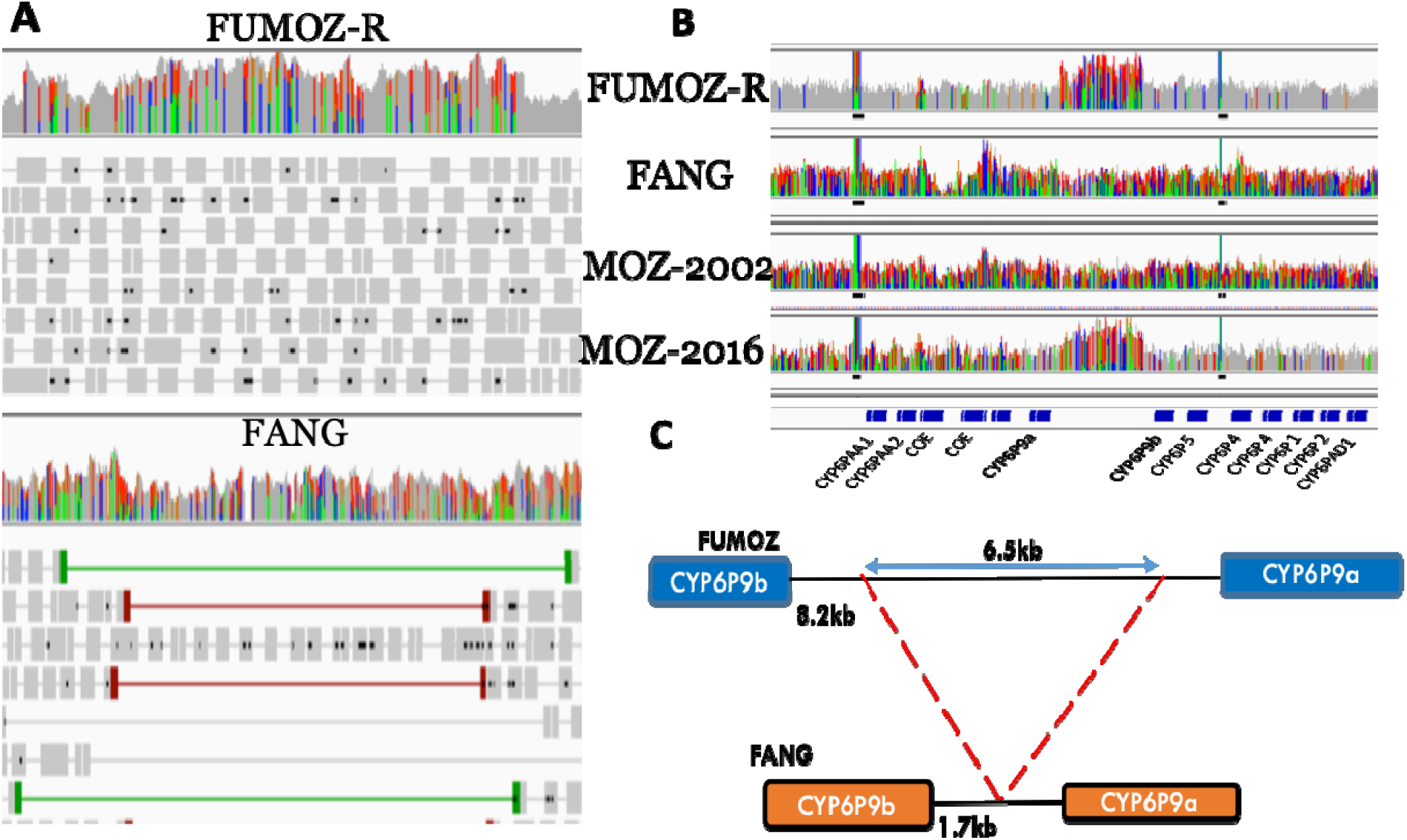
Detection of the 6.5kb intergenic insertion between *CYP6P9a* and *CYP6P9b* in *An. funestus* mosquitoes. (A) Screenshot from the integrative genomics viewer (IGV), showing the coverage depth and aligned reads for the pyrethroid resistant FUMOZ-R strain (upper) and the fully susceptible FANG (lower) using the pooled template whole genome sequence alignments (Pool-seq). The coverage depth plots show deeper coverage in this region in FUMOZ-R but not in FANG. The FANG alignment contains read pairs with unusually long insert sizes, indicated in red in the lower panel (thick lines represent reads, read pairs are linked by thin lines). (B) IGV screenshot showing an increase coverage depth of reads between *CYP6P9a* and *CYP6P9b* in FUMOZ-R with a loss of polymorphism (grey) compared to the susceptible FANG strain using Pool-seq data. The same contrast is observed between the 2002 sample from Mozambique before LLIN scale up (MOZ-2002) (high diversity and low coverage of read depth) 2016 sample (MOZ-2016) where a greater coverage depth is observed at the intergenic region corresponding to the insertion of the 6.5kb fragment. (C) Schematic representation of the 6.5kb insertion in FUMOZ-R in comparison to FANG.

The results also indicate that the 6.5 kb insertion between *CYP6P9a* and *CYP6P9b* was present only in southern Africa population of Malawi, where it was nearly fixed (only a single read in Malawi supported the deletion haplotype) (Table S1). However, populations from other parts of Africa showed no evidence of the insertion haplotype. Evidence that the insertion existed (albeit at low frequency) in the early 2000s comes from its presence in the FUMOZ-R colony, which was colonized from the field in Mozambique in 2000, and subsequently selected for insecticide resistance, which appears to have fixed the insertion haplotype in colony.

### Investigating the genomic origin of the 6.5kb insertion

To identify the genomic origin of the inserted 6.5 kb sequence, its entire sequence was used to search the *An. funestus* FUMOZ AfunF3 reference genome assembly using BLASTn implemented in the VectorBase web resource. The results indicated that the sequence occurred at another different location in the genome, on scaffold CM012071.1 (S2A Fig). That location is between 8,296,288 and 8,555,956, approximately 260 kb away from the CYP6 cluster on the same scaffold (therefore, on the same chromosome 2). Both locus are on the chromosome 2R in the same genomic region as the P450s cluster rp1 QTL [17] which confers high pyrethroid resistance and shown to have undergone a major selective sweep. In addition, short (100 bp) sequences from the left and right ends of the insertion were used to conduct BLASTn searches of AfunF3 and confirmed the results obtained with the full-length insertion sequence. Finally, clipped sequences from immediately to the left and right of the insertion were used to conduct BLASTn searches of AfunF3. The results (matches only adjacent to the CM012071.1:8,296,288-8,555,956 region) confirmed that the “parent” sequence of the insertion between *CYP6P9a* and *CYP6P9b* came from CM012071.1:8,296,288-8,555,956. This putative genomic “parent” sequence of the insert contains no annotated protein coding genes but there is a large assembly gap in the region. The orthologous region in the *Anopheles gambiae* genome is also on chromosome arm 2R (S2B Fig). The protein coding genes flanking the insertion sequence, AFUN008344 and AFUN008346, are orthologous to AGAP002842 and AGAP002845, respectively. *An. gambiae* also have no annotated protein coding genes between AGAP002842 and AGAP002845, suggesting that *An. funestus* may not have also. Two micro-RNAs annotated in both species are outside of the insertion sequence. Despite the lack of annotated genes, the region is transcribed as supported by FUMOZ-R RNAseq showing a large transcribed region (S2C Fig), with some evidence of splicing, covering the two annotated micro-RNAs. Whether this transcript is processed to form mature micro-RNAs is not known.

### Comparative promoter analysis of the *CYP6P9a*-*CYP6P9b* intergenic region

The location of the 6.5kb insertion in the intergenic region between *CYP6P9a* and *CYP6P9b* (Fig 1C) indicated that this insertion could impact the regulation of these genes. This is strongly supported by their high expression in FUMOZ-R and Southern Africa mosquito populations. To verify this hypothesis, the full length intergenic fragment (8.2kb) was successfully PCR amplified and cloned (S2D Fig) and used for a luciferase reporter assay (Pgl3 FZ6P9a8.2) comparatively with three other construct vectors without the 6.5kb SV (Pgl3 FZ6P9a0.8, Pgl3 FG6P9a0.8, and Pgl3 LRIM). These constructs were transfected alongside Renilla luciferase plasmid in *An. gambiae* 4a-2 cell line. The samples were lysed and luciferase activity was measured and normalized against Renilla activity. The luciferase activity of the construct with the 6.5 kb SV from FUMOZ-R, Pgl3 FZ6P9a8.2, was 5-fold significantly higher (P=0.00012) than that of the same strain without it, FZ6P9a0.8 and 17.5-fold higher than the FANG fragment, Pgl3 FG6P9a0.8 (P<0.0001). This result implies that this insertion likely contains cis-regulatory elements acting as gene regulation enhancers and driving the high expression of *CYP6P9a* and *CYP6P9b* observed (Fig 2A).

**Figure 2:**
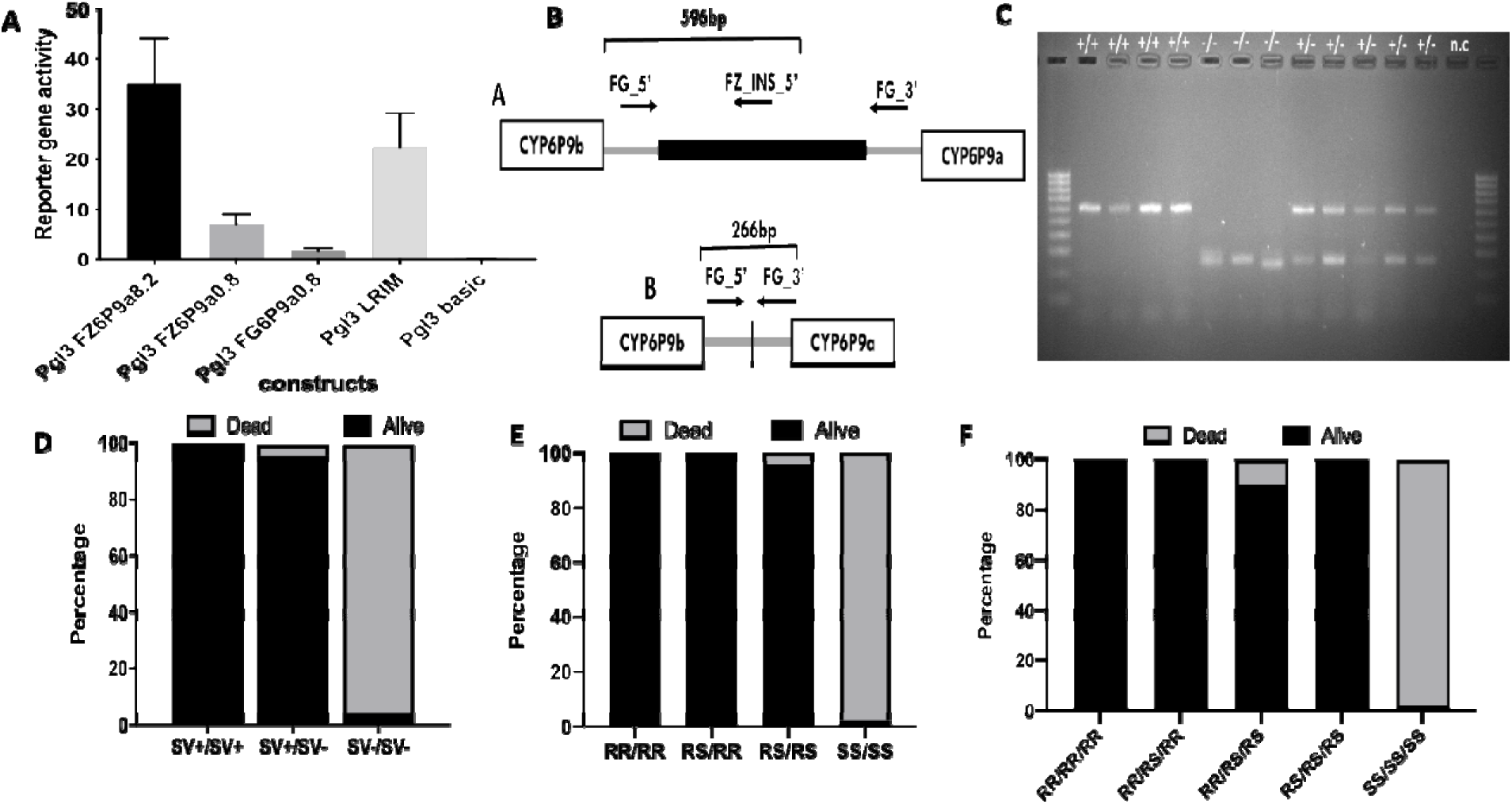
Functional characterization of the 6.5 kb SV and design of a PCR diagnostic assay. (A) Comparative Luciferase reporter assay (normalised to renilla fluorescence) of various intergenic fragments between *CYP6P9a* and *CYP6P9b*. Pgl3 FZ6P9a8.2 is the fragment for the full 8.2 intergenic region containing the 6.5kb insertion showing a very elevated activity compared to others. Pgl3 FZ6P9a0.8 is the 800 bp region upstream of *CYP6P9a* containing cis-regulatory elements used to design PCR-assay for *CYP6P9a* [13] whereas Pgl3 FG6P9a0.8 is the FANG version. Error bars show +/- SD. (B) Schematic representation of the design of the PCR assay to detect the 6.5 kb insertion using three primers two located immediately outside of the SV (FG_5’ and FG_3’; size of 266bp if no SV) and one located within the 6.5kb SV allowing to detect its presence (expected band 596bp). (C) Agarose gel showing the three genotypes indicating the presence (+) or absence (-) of the insertion using F_8_ FUMOZ-R x FANG crosses. Ladder: 100bp, n.c.: negative control. (D) Distribution of the 6.5 kb *SV* genotypes between susceptible (Dead) and resistant (Alive) mosquitoes after WHO bioassays with 0.75% permethrin showing a very strong correlation between the 6.5 kb SV and pyrethroid resistance phenotype. (E) Distribution of the combined genotypes of 6.5 kb SV and that of *CYP6P9a* after WHO bioassays with 0.75% permethrin showing that genotypes of both loci combined to increase the pyrethroid resistance. (F) Distribution of the combined triple genotypes of 6.5kb besides that of both P450s *CYP6P9a* and *CYP6P9b* after WHO bioassays with 0.75% permethrin showing that the triple resistance genotypes combined to further increase the pyrethroid resistance

### A simple PCR assay to detect 6.5 kb structural variant and associated pyrethroid resistance

The 8.2kb (SV+) and 1.7 kb (SV-) intergenic regions were used to design a simple PCR assay capable of detect the 6.5kb structural variant. The PCR is comprised of 3 primers, two in the flanking region and one in the 6.5kb region (Fig 2B). The presence of the SV is shown by the amplification of a fragment of 596 bp (SV+) whereas the absence is shown by a band at 266 bp (SV-) (Fig 2C). To assess the efficacy of this novel PCR, we genotyped the FANG-S and FUMOZ-R strains for the 6.5 kb structural variant and found that all FUMOZ-R mosquitoes genotyped were homozygous for insertion (SV+/SV+) while all the FANG where homozygous without insertion (SV-/SV-).

To assess the association between this 6.5kb SV and pyrethroid resistance in *An. funestus*, F_8_ mosquitoes obtained from crossing FANG and FUMOZ-R which had been exposed to permethrin 0.75% for 180 minutes to get highly resistant mosquitoes and 30 minutes to get highly susceptible mosquitoes were genotyped for this SV. The results revealed that those alive after 180 minutes exposure were mainly homozygote (21/45) and heterozygote (22/45) for the 6.5kb SV. Only 2/45 alive were lacking the 6.5kb SV. Genotyping of the 41 highly susceptible mosquitoes, 40/41 were homozygote without the 6.5kb SV and 1/41 heterozygote (Fig 2D). Consequently, a strong association was found between permethrin resistance and 6.5kb SV genotype with a highly significant odds ratio when comparing the homozygote (SV+/SV+) to the homozygote (SV-/SV-) (SS) (OR=2079.4, *P*<0.001, Fischer’s exact test). Similarly, a significantly higher ability to survive was observed for heterozygote (SV+/SV-) when compared to homozygote (SV-/SV-) (OR=600.25, *P*<0.001, Fischer’s exact test) (Fig 2D) (S2 Table). Moreover, analysis of the combined effect of the 6.5kb SV with the two nearby P450s revealed an increased survivorship when the 6.5kb SV is combined to either *CYP6P9a* (Fig 2E: S2 Table) or *CYP6P9b* (S2 Table) with an even greater chance to survive when a mosquito is triple homozygotes for the resistant allele at the three loci (Fig 2F: S2 Table).

### Enhancer’s effect of the 6.5kb structural variant on the expression of nearby genes

To confirm the potential enhancer role of the 6.5kb SV on the regulation of nearby P450 genes as suggested by the luciferase promoter assay, quantitative real time PCR was used to compare the expression of the nearby *CYP6P9a* and *CYP6P9b* P450 genes. Mosquitoes obtained from F_2_ crossing of FUMOZ-R and FANG were pooled into three groups based on their genotypes (SV+/SV+, SV+/SV- and SV-/SV-) for the 6.5kb SV (Fig 3A and 3B). Analysis revealed that the mosquitoes possessing the 6.5 kb SV (SV+/SV+ and SV+/SV) had a significantly higher expression level for *CYP6P9*a and *CYP6P9b* as opposed to those without the 6.5kb SV (SV-/SV-) (Fig 3C). Expression level of *CYP6P9*a for mosquitoes with SV+/SV+ genotype was about 1.7-fold (P=0.03) higher than that of SV+/SV- heterozygote genotype and 5.2-fold higher than the homozygote without SV (SV- /SV-) (P=0.005). A similar pattern was observed for *CYP6P9b* with homozygote SV+/SV+ genotype expressing *CYP6P9b* about 2-fold higher than heterozygote mosquitoes (P=0.003) and 4-fold higher than the homozygote SV-/SV- (P=0.001). This suggests a strong association between the 6.5kb SV and increased expression of *CYP6P9a* and *CYP6P9b* likely as a result of the presence of cis-regulatory elements in the 6.5kb acting as enhancers.

**Figure 3:**
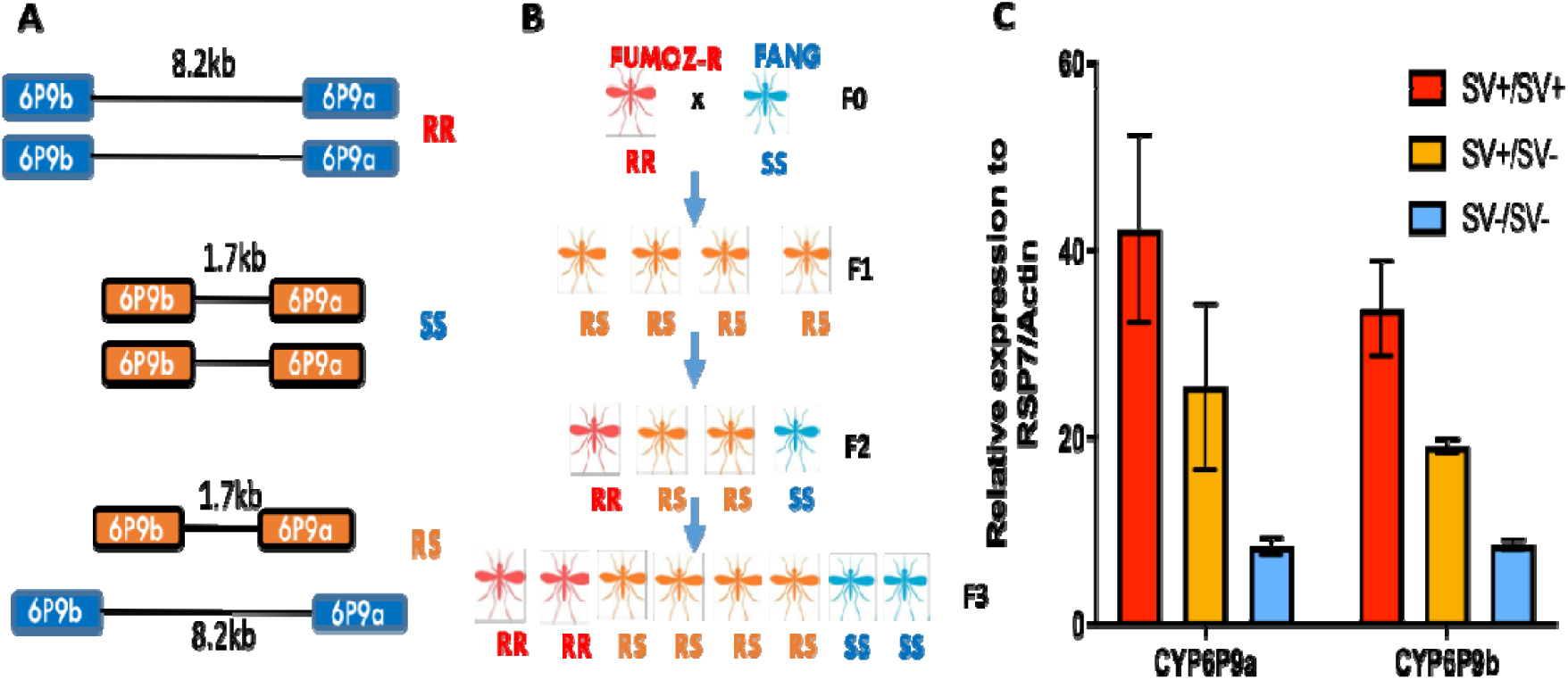
Investigation of the enhancer role of the 6.5kb SV using comparative gene expression profile of the three SV genotypes on *CYP6P9a* and *CYP6P9b*: (A) Schematic representation of the three 6.5kb SV genotypes expected in *An. funestus* mosquitoes during the crossing process. (B) Experimental design of the crossing between the pyrethroid resistant strain FUMOZ-R (with 100% SV) and the fully susceptible FANG (no SV) up to the F_3_ generation. Expected segregation of genotypes is shown. (C) Differential qRT-PCR expression for different insertion genotypes of two cytochrome P450 genes at the immediate vicinity of the 6.5 kb SV. Error bars represent standard deviation (n = 3). P<0.05 is represented by *, ** for p< 0.01, *** for p<0.001.

### Geographic distribution of 6.5kb structural variant across Africa

This novel diagnostic PCR was tested on field mosquitoes collected from various African regions to determine the spread of this insertion across the continent. Genotyping of the 6.5kb SV was successful in all the 7 countries revealing that this structural variant is present in Southern Africa mosquito populations while absent from those collected in the Central and Western Africa (Fig 4A). The frequency of 6.5 kb SV was very high in southern Africa ranging from 82% in Zambia, 92% in Malawi and 100% in Mozambique (Fig 4B). Tanzania and Eastern Democratic Republic of Congo presented intermediate frequencies (43.94%) and (72.2%) respectively. The 6.5kb SV was completely absent in Western (Ghana), Eastern (Kenya, Uganda) and Central Africa (Cameroon and DRC-Kinshasa). Hence this structural variant is confined to Southern Africa like the *CYP6P9a* and *CYP9P9b* P450 based pyrethroid resistance [13, 14] and contrary to the L119F and A296S resistance-to-dieldrin (RDL) mutation, which confer DDT and dieldrin resistance respectively [9]. Samples from Tanzania and DRC-Mikalayi that show a segregation of different genotypes, were further used to assess the potential association between the 6.5kb SV, *CYP6P9a* and *b* (Fig 4C; S3A and S3B Fig). In Tanzania, a greater linkage of the three markers was observed with 54.8% (17/31) identical genotypes detected while the value was lower in DRC-Mikalayi with 32% (8/25). Considering only *CYP6P9a* and the 6.5kb SV, 12% (3/25) and 16% (5/31) had the same genotype for both markers for DRC-Mikalayi and Tanzania respectively. A low genotypic frequency of 8% (2/25) was obtained for samples with the same genotype for *CYP6P9b* and the 6.5kb SV for DRC-Mikalayi. With regard to *CYP6P9a* and *CYP6P9b*, 48% (12/25) and 25.81% (8/31) from DRC-Mikalayi and Tanzania respectively had the same genotype (Fig 4C).

**Figure 4:**
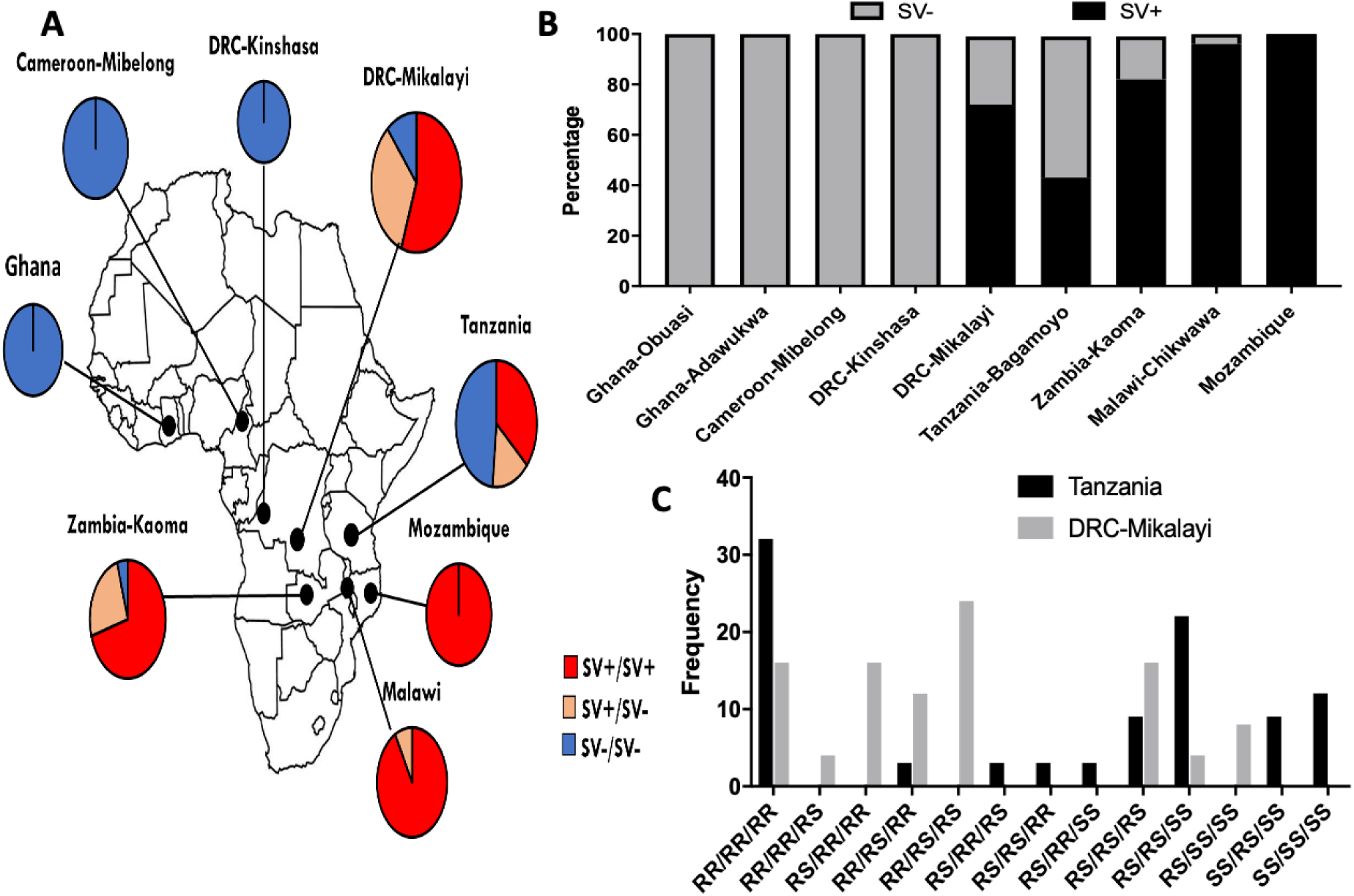
Geographical distribution of the 6.5 kb SV across Africa; (A) Map of Africa showing the genotypic distribution of the 6.5kb SV across the continent which correlated with the *CYP6P9a_R* and *CYP6P9b_R* allele distribution. (B) Histogram showing the frequency of the SV+ and SV-alleles across Africa revealing a clear gradient from Mozambique (100% SV+) to West/Central Africa (0% SV+). (C) Distribution of the combined genotypes *CYP6P9a, CYP6P9b* and 6.5kb SV in Tanzania and DRC Mikalayi showing and extensive segregation of the genotypes to these loci in the field.

### 6.5kb Structural variant is associated with reduced bed net efficacy

The association between the 6.5 kb SV and the efficacy of bed net was assessed using mosquitoes previously used to validate *CYP6P9a* [13] and *CYP6P9b* [14]. Briefly, mosquitoes were obtained from crossing the susceptible FANG laboratory strain (where the 6.5kb is totally absent) and highly pyrethroid resistant laboratory FUMOZ-R (where the 6.5kb SV is fixed). Mosquitoes from the crossing were reared to the F_6_, which was used for cone assays and to perform a release recapture in experimental huts.

#### Cone assays

To validate the ability of the 6.5kb SV marker to predict the impact of resistance on the efficacy of LLINs, we genotyped F_6_ samples obtained from cone assay with PermaNet 2.0.and PermaNet 3.0 (side). The mortality rates were 31.1% and 40.7% for PermaNet 2.0.and PermaNet 3.0 respectively with no significant difference (*P* value = 0.380). With PermaNet 2.0, a significant difference in the distribution of genotypes of the SV was observed between dead and alive mosquitoes (Chi^2^=892; P<0.0001, Chi-square) (Fig 5A). Comparing the proportion of each SV genotype between alive and dead mosquitoes revealed that SV+/SV+ homozygote mosquitoes were significantly more likely to survive exposure to PermaNet 2.0 than those completely lacking the SV (SV-/SV-) (OR=1798.3; CI=97.6-33141; P<0.0001, Fisher’s exact test). Heterozygote mosquitoes were also more able to survive exposure to PermaNet 2.0 than homozygotes lacking the SV (SV-/SV-) (OR=261.5; CI=15.4-4447.3; P<0.0001, Fisher’s exact test). SV+/SV+ homozygote mosquitoes were also more able to survive than heterozygotes (OR=7.4; CI=2.5-21.5; P=0.0002, Fisher’s exact test) showing an additive effect of the 6.5kb SV on the resistance phenotype. When comparing at the allelic level, it was observed that possessing a single SV+ allele provides a significant ability to survive exposure to PermaNet 2.0 compared to the SV-allele (OR=27.7; CI=12.9-59.0; P<0.0001, Fisher’s exact test) (Fig 5A; S3 Table). A similar trend was observed with PermaNet 3.0 (side) (Fig 5A; S4 Table) whereas we could not assess PermaNet 3.0 (Top) as there was no survivor after exposure.

**Figure 5:**
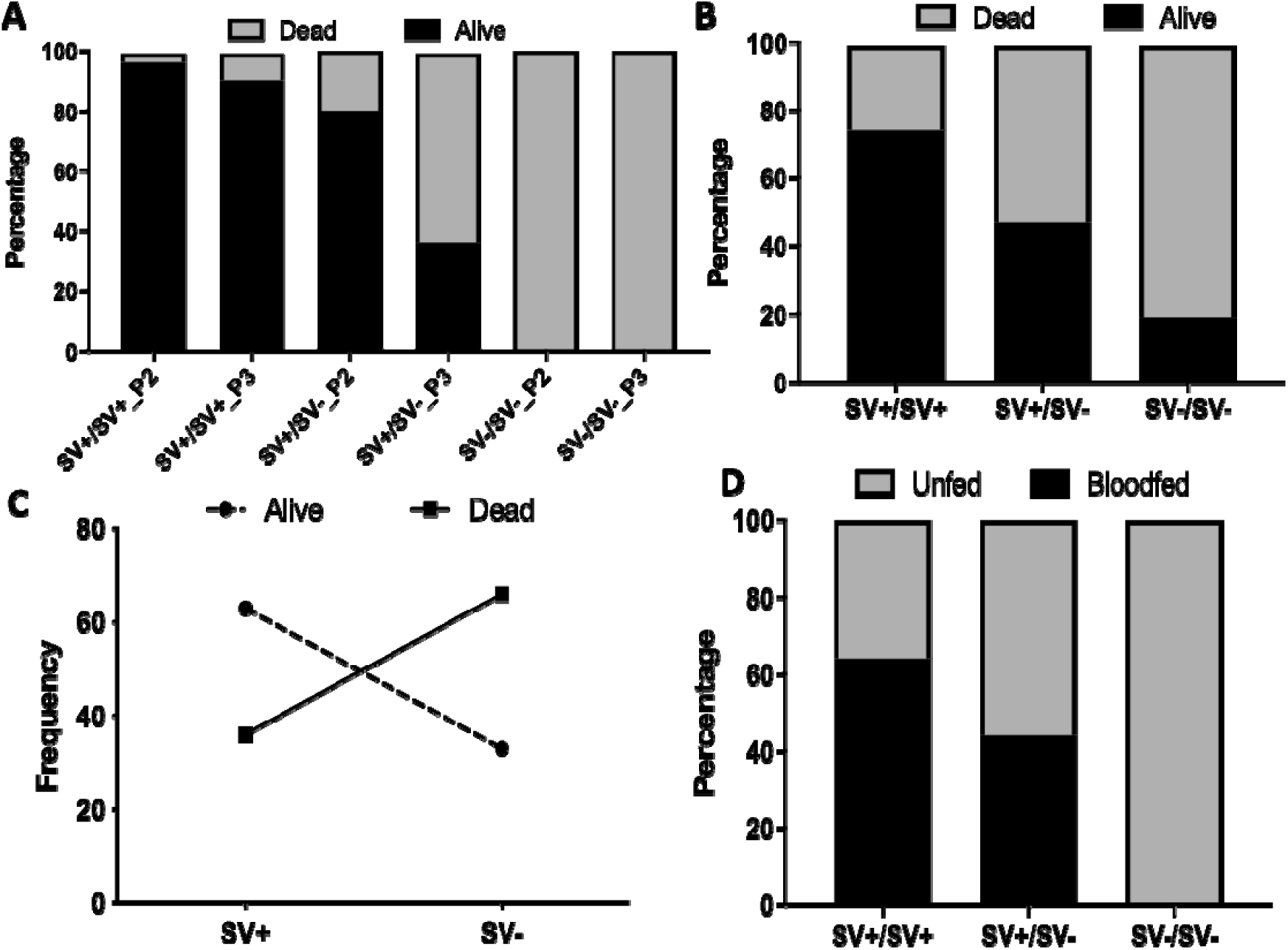
Impact of the 6.5 kb enhancer on the efficacy of insecticide-treated nets using experimental hut trials: (A) Distribution of 6.5kb SV genotypes among mosquitoes that survive exposure to PermaNet 2.0 (P2) and PermaNet 3.0 (side) (P3) and those that died using cone assays showing an increase ability to withstand exposure to bed net when possessing one and two copies of the 6.5 kb SV. (B) Distribution of 6.5 kb *SV* genotypes between dead and alive mosquitoes after exposure to PermaNet 2.0 net in huts showing that the 6.5 kb SV significantly allows mosquitoes to survive exposure to this bed net. (C) Correlation between frequency of 6.5 kb SV alleles and ability to survive exposure to PermaNet 2.0. (D) Distribution of the 6.5 kb *SV* genotypes between blood fed and unfed mosquitoes after exposure to the PBO-based net PermaNet 3.0 in huts showing that SV+ allele increases the ability to take a blood meal even for PBO-based nets.

#### Experimental hut trial

Mosquitoes previously collected from experimental huts after 4 consecutive nights of release-recapture and used to validate the *CYP6P9a* and *CYP6P9b* were analysed for the 6.5kb SV. The mortality rate were 98.7% for PermaNet 3.0 and 33.3% for PermaNet 2.0 [13]. To avoid the bias due to exophily and feeding status, only samples that were unfed and in the room were used. A significant difference was observed in the distribution of genotypes of the SV among the alive and dead mosquitoes (Chi^2^=40.2; P<0.0001, Chi-square) (Fig 5B and 5C).Comparison of the proportion of different genotypes among the dead and alive revealed that 6.5kb SV homozygote (SV+/SV+) were able to significantly survived exposure to PermaNet 2.0 more than those without the 6.5kb SV (SV- SV-) [OR =27.7; CI: 13.0-59.0; P < 0.0001, Fisher’s exact test] (Table 1). Mosquitoes heterozygote for the 6.5kb SV (SV+/SV-) also survived exposure to PermaNet 2.0 more than those homozygote without the 6.5 SV (SS) [OR = 7.5; CI:3.8-14.8; P < 0.0001, Fisher’s exact test]. However, the 6.5kb SV heterozygote (SV+/SV-) survived less than the 6.5kb SV homozygote (SV+/SV+) (OR =3.7; CI = 1.9-7.1; P =0.0001, Fisher’s exact test) suggesting that there is an additive effect associated with possessing 2 copies rather than 1. This was further demonstrated by the ability of mosquitoes having a single 6.5kb structural variant to survive more than those without (Fig 5B). A similar trend was observed when the analysis was performed without excluding the exophilic status and blood-fed (S5 Table) as well as for PermaNet 3.0 (S4A Fig; S6 and S7 Table).

**Table 1:**
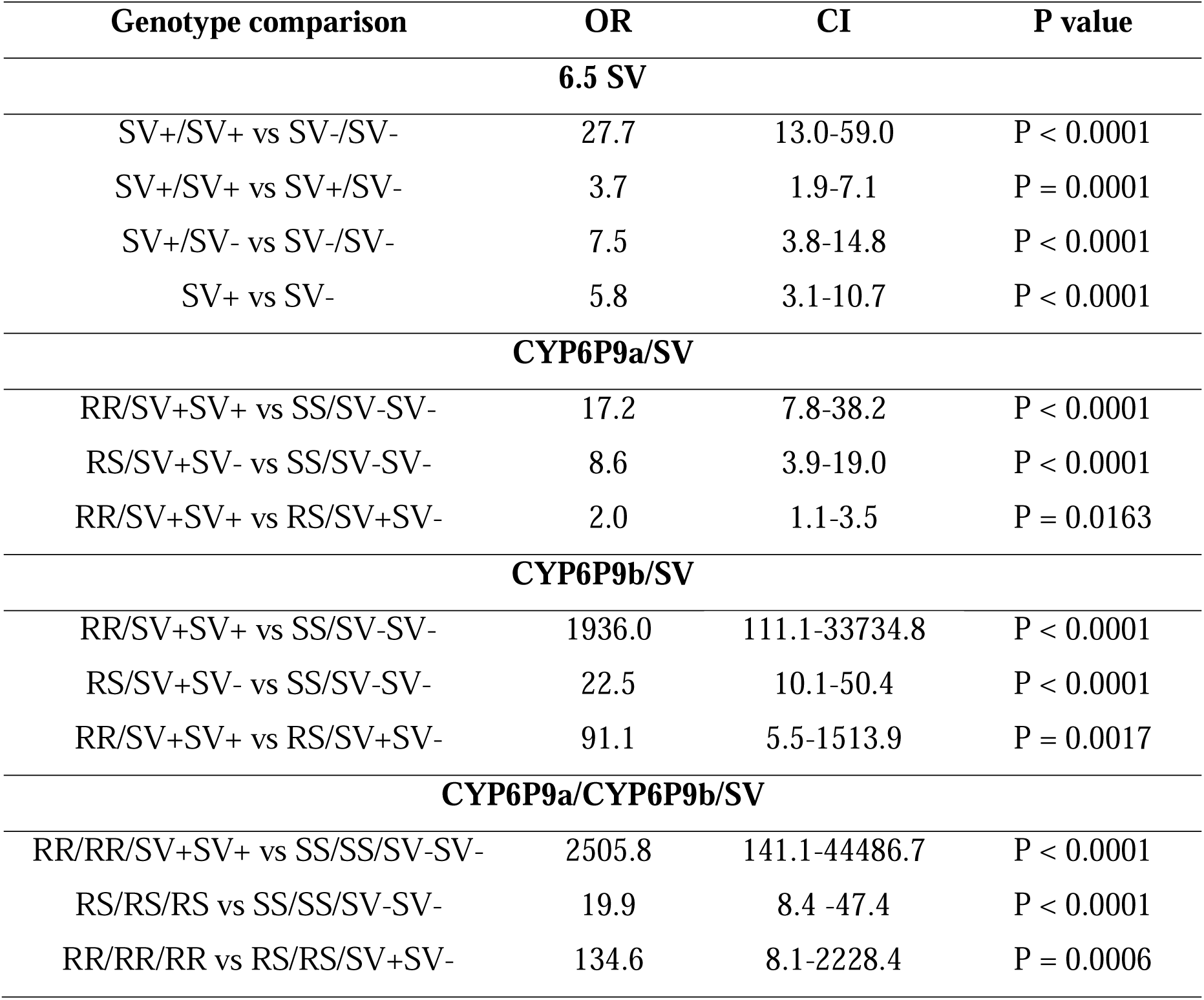
Correlation between genotypes of the 6.5kb SV and ability to survive exposure to PermaNet 2.0 in experimental huts using unfed samples.

The distribution of the 6.5kb SV in blood-fed mosquitoes was assessed to determine any association between this structural variant and the ability of mosquitoes to blood feed in the presence of the bed-net. A greater number of mosquitoes were able to blood-fed with PermaNet 2.0 (14.8 %) compared to PermaNet 3.0 (6.8%) [13]. A significant difference was observed for the distribution of the 6.5kb SV genotypes between blood-fed and unfed collected from huts with both PermaNet 3.0 (chi square=18.4; P<0.0001) and PermaNet 2.0 (chi square= 60.2; P<0.0001). A significant association was observed between the 6.5kb SV genotypes and the ability of mosquitoes to blood feed when exposed to PermaNet 3.0. Mosquitoes homozygote for the 6.5kb SV (SV+/SV+) were more likely to blood feed in the presence of PermaNet 3.0 than those homozygote without (SV-/SV-) (OR=355.2; CI=21.4-5889.4; P < 0.0001) and heterozygote for the 6.5kb SV (SV+/SV-) (OR=2.6; CI=1.3-4.0; P = 0.00489). Mosquitoes heterozygote for the 6.5 SV (SV+/SV-) were also able to blood feed more that those without the structural variant (SV-/SV-) (OR=158.3; CI=9.6-2620.0; P = 0.0004) (Fig. 5D; Table 2). Analysis of genotype distribution for PermaNet 2.0 gave similar trends but these were not significant apart from SV+/SV- vs SV-/SV- (S4B Fig, S8 Table).

**Table 2:** Correlation between genotypes of the 6.5kb SV and ability to blood-feed when exposed to PermaNet 3.0 in experimental huts.

| Genotype comparison | OR | CI | P value |
| --- | --- | --- | --- |
| **6.5 SV** |  |  |  |
| SV+/SV+ vs SV-/SV- | 355.2 | 21.4-5889.4 | P < 0.0001 |
| SV+/SV+ vs SV+/SV- | 2.6 | 1.3-4.0 | P = 0.0048 |
| SV+/SV- vs SV-/SV- | 158.3 | 9.6-2620.0 | P = 0.0004 |
| SV+ vs SV- | 2.5 | 1.4-4.4 | P = 0.0020 |
| **CYP6P9a/SV** |  |  |  |
| RR/SV+SV+ vs SS/SV-SV- | 4.5 | 2.5-8.2 | P < 0.0001 |
| RS/SV+SV- vs SS/SV-SV- | 2.0 | 1.1-3.6 | P = 0.0205 |
| RR/SV+SV+ vs RS/SV+SV- | 2.3 | 1.3-4.0 | P = 0.0047 |
| **CYP6P9b/SV** |  |  |  |
| RR/SV+SV+ vs SS/SV-SV- | 4.5 | 2.5-8.3 | P < 0.0001 |
| RS/SV+SV- vs SS/SV-SV- | 2.3 | 1.3-4.1 | P = 0.0062 |
| RR/SV+SV+ vs RS/SV+SV- | 2.0 | 1.1-3.6 | P = 0.0159 |
| **CYP6P9a/CYP6P9b/ SV** |  |  |  |
| RR/RR/SV+SV+ vs SS/SS/SV-SV- | 5.0 | 2.7-9.0 | P < 0.0001 |
| RS/RS/RS vs SS/SS/SV-SV- | 2.1 | 1.2-3.7 | P = 0.0141 |
| RR/RR/RR vs RS/RS/SV+SV- | 2.4 | 1.3-4.3 | P = 0.0029 |

### 6.5kb Structural variant combines with *CYP6P9a* and *CYP6P9b* to further reduce LLIN efficacy

We next assessed the impact of different combinations of genotypes for the 3 loci on the efficacy of LLINs. The cone assays results revealed an increased survivorship when the 6.5kb SV is combined to *CYP6P9a* for PermaNet 2.0 (S5A Fig; S3 Table) and PermaNet 3.0 (side) (S5B Fig; S4 Table). Similar patterns were observed when combined to *CYP6P9b* (S5C and S5D Fig; S3 and S4 Table). Mosquitoes with triple homozygotes for the resistant allele at the three loci exhibited a greater ability to survive than all other genotype combinations for both nets (S5E and S5F Fig; S3 and S4 Table).

Experimental hut trials also showed that triple homozygote RR/RR/SV+SV+ were more likely to survive exposure to PermaNet 2.0 than any other combined genotype (Fig 6A; Table 1). The RR/RR/SV+SV+ survived more than the SS/SS/SV-SV- (OR=2505.8; CI: 141.1-44486.7; P<0.0001), RR/RS/SV+SV- (OR=31.0; CI=1.8 to 529.3; P = 0.0177) and RS/RS/SV+SV- (OR= 134.6CI: 8.1-2228.4; P = 0.0006). There was no significant difference between the RR/RR/SV+SV+ and RR/RR/SV+SV- genotypes (OR= 1; CI= 0.02 to 50.9; P= 1.0) and both had a similar trend in terms of survival with the same odds ratio (Fig 6B; Table 1). Hence, there is an additive advantage associated with the mosquito combined genotype at the 3 loci which confers higher level of pyrethroid resistance leading to a greater reduction in bednet efficacy. The impact decreased in the order RR/RR/SV+SV+=RR/RR/SV+SV- >RR/RS/SV+SV-> RS/RS/SV+SV-> SS/SS/SV-SV-. This additive effect was also observed for the PBO-based PermaNet 3.0 net although at a lower extent (OR=423; P<0.0001; RRR/RR/SV+/SV+ vs SS/SS/SV-/SV-)(S4C Fig; S6 and S7 Table). Moreover, an increased ability to survive exposure to LLIN is even observed when combining genotypes of the 6.5kb SV only to either of *CYP6P9a* (S6A and S6B Fig) or *CYP6P9b* (S7A and S7B Fig).

**Figure 6:**
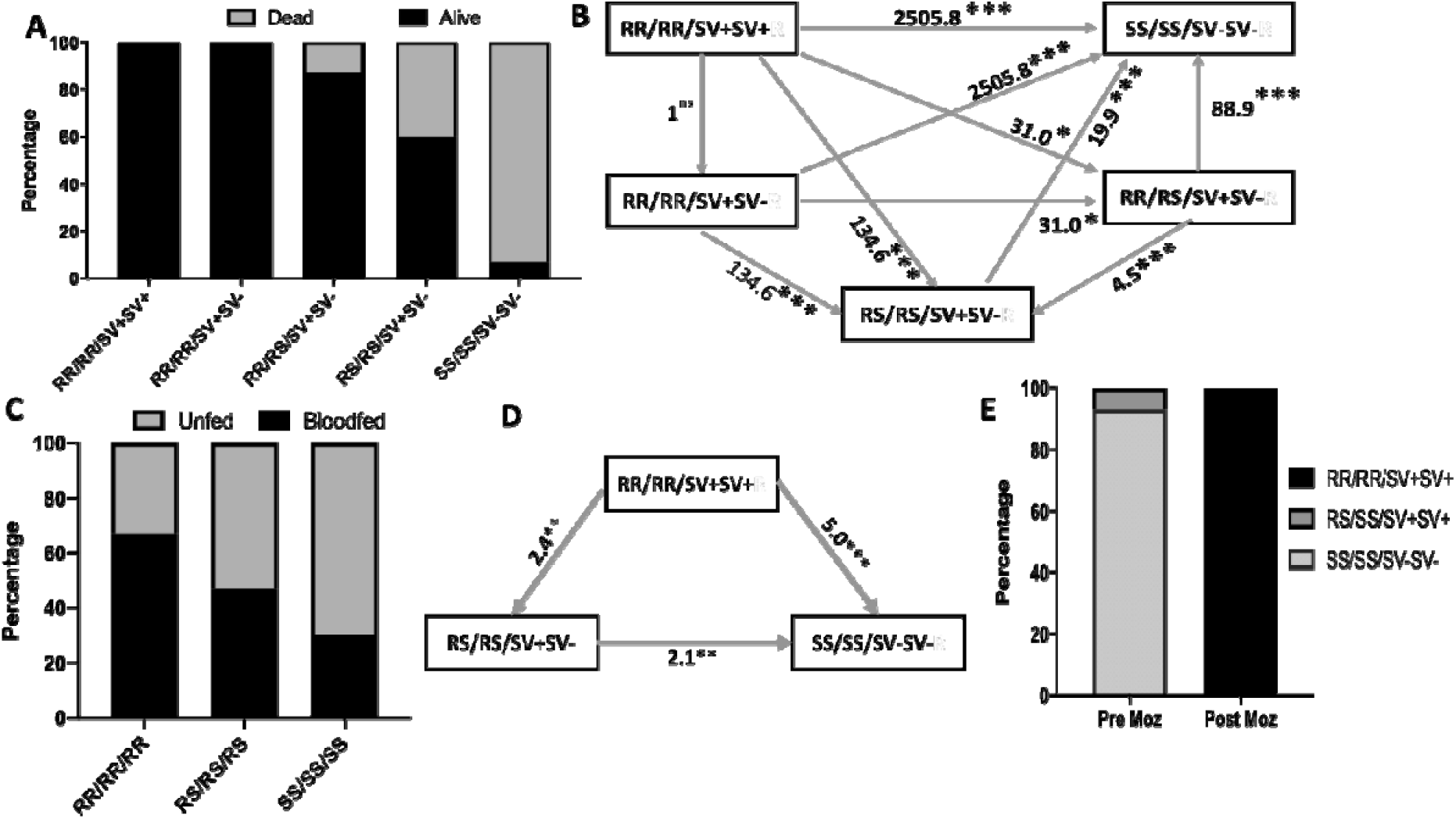
The 6.5 kb *SV* combines with both *CYP6P9a*_R and *CYP6P9b_R* P450 alleles to further reduce the efficacy of insecticide-treated nets: (A) Distribution of combined genotypes for the 6.5 kb SV *CYP6P9a* and *CYP6P9b* showing that three markers combine to increase survivorship in the presence of PermaNet 2.0. (B) Ability to survive exposure to PermaNet 2.0 for the various combined genotypes for the 6.5 kb SV *CYP6P9a* and *CYP6P9b*. *P < 0.05, **P < 0.01, ***P < 0.001. (C) Distribution of the combined genotypes for the 6.5 kb SV, *CYP6P9a* and *CYP6P9b* in mosquitoes collected after exposure to PermaNet 3.0 in huts show that the 3 loci combine to increase blood feeding ability. (D) The triple RR/RR/SV+SV+ homozygote mosquitoes for the 6.5 kb SV, *CYP6P9a* and *CYP6P9b* exhibit a greater blood feeding ability than other genotypes. (E) Impact of bed net scale up in Mozambique on selection of the 6.5 kb SV, *CYP6P9a* and *CYP6P9b* genotypes with no triple homozygote genotype before (Pre Moz) and 100% after (Post Moz) scale-up of insecticide-treated nets

In addition to the ability to survive bednet exposure, RR/RR/SV+SV+ also had a higher ability to blood feed compared to SS/SS/SV-SV- (OR=5.0; CI: 2.7-9.0; P < 0.0001) and the other combinations in the presence of PermaNet 3.0 (Fig 6C and 6D). This additive effect was also observed when combining genotypes of the 6.5kb SV only to either of *CYP6P9a* (S6C and S6D Fig) or *CYP6P9b* (S7C and S7D Fig). For PermaNet 2.0 no significant association was observed (S4D Fig; S8 Table).

### Scale-up of bednets is driving combined selection of 6.5kb SV, *CYP6P9a* and *CYP6P9b*

We assessed how these three loci are being selected in the field by insecticide-based interventions using mosquito samples (27) dating before (2002) and after scale-up of bed nets in southern Africa (2016) more precisely in Mozambique. The results revealed that before the introduction of bed nets the distribution of genotypes were skewed towards the SS/SS/SS (92.59%) and RS/SS/SS (11.76%) combined genotype than the RR/RR/RR (0%). Genotyping of the mosquitoes collected after scale-up of bed nets showed that the distribution of the combined genotype frequencies had completely changed with the RR/RR/RR becoming fixed among mosquitoes in 2016 (100%) (Fig 6E and S8 Fig).

## Discussion

This study investigated the role that structural variations in cis-regulatory regions play in cytochrome P450-mediated metabolic resistance to pyrethroids in mosquitoes by focusing on a 6.5kb intergenic insertion between two duplicated P450 in the malaria vector *An. funestus*. This work showed that the 6.5kb SV acts an enhancer for nearby duplicated P450 genes *CYP6P9a* and *CYP6P9b* leading to their increased overexpression. This 6.5kb SV is strongly associated with an aggravation of pyrethroid resistance which reduces the efficacy of pyrethroid-only nets but also to some extent that of PBO-nets. This study improves our understanding of the molecular processes that drive P450-based resistance to insecticides in mosquitoes while providing an additional marker for monitoring pyrethroid resistance in wild *An. funestus* populations.

### 1-The 6.5kb SV acts as an enhancer for the up-regulation of duplicated P450 genes

The whole genome Pool-seq confirmation of the 6.5kb insertion in field populations of *An. funestus* supported the previous report in the lab pyrethroid resistant FUMOZ-R strain [13]. This study has revealed that this insertion acts as an enhancer to the over-expression of both duplicated P450 genes *CYP6P9a* and *CYP6P9b* which are key pyrethroid resistance genes in FUMOZ-R [17, 18] and in southern Africa [13, 19]. The first line of evidence that the 6.5kb insertion serves as an enhancer was provided by the comparative promoter assays performed between the promoter with the 6.5kb SV and the core promoter region of *CYP6P9a*. The greater promoter activity observed in the presence of the SV than in the absence shows that this insertion increases the over-expression of these P450 genes. This is further supported by the previous report that the 6.5kb is highly enriched with regulatory elements including several transcription factors binding sites as well as several TATA (35), CCAAT (12) and GC (11) sequences. Furthermore, the presence of 51 sites for the Cap n Collar C (CnCC) and the Muscle aponeurosis fibromatosis (Maf) transcription factors sites which are known xenobiotic sensors in insects [13, 16] supports the role likely played by this insertion in enhancing the regulation of neighbouring genes. The analysis of the composition of the 6.5kb insertion provides a good insight into the architectural characteristics of enhancers in mosquitoes which is similar to that of mammalians which have also be shown to include the multiple transcription factor binding sites as well as transcriptionally activating and repressing domains in the same enhancer [20].

Secondly, at the transcriptional level, the relative expression of *CYP6P9a* and *CYP96P9b* were shown to correlate with the 6.5kb SV genotypes. The presence of this SV is strongly associated with increased expression of these two genes (SV+/SV+>SV+/SV->SV- /SV-) which are located upstream and downstream of this SV. It is well documented that insertions by introducing novel cis-acting elements into the regulatory regions can either alter expression of a gene or disrupt it [21]. The fact that this 6.5kb is enhancing the expression of two genes is in line with what is already known about enhancers which have be shown to be able to regulate genes in cis-position in either upstream (e.g *CYP6P9a*) or downstream (e.g *CYP6P9b*) or even in intron [22]. The fact that the presence of the 6.5kb SV impacts the two genes is also in line with previous reports stating that enhancers can regulate the expression of multiple genes [22]. It will be good to establish whether this 6.5kb regulates the expression of other genes besides *CYP6P9a/b* notably on the cluster of P450s in this *rp1* resistance locus [17].

Thirdly, the geographical distribution of the 6.5kb SV tightly correlated with the high over-expression of *CYP6P9a* and *CYP6P9b* in the FUMOZ-R strain and across southern African countries [13, 19] but a low expression elsewhere in Africa where this SV is absent such as in Central Africa, East and West Africa [23]. This observation further supports that the 6.5kb is driving the up-regulation of both genes. This is similar to what was reported in *Drosophila melanogaster* where over-expression of the cytochrome P450 gene *Cyp6g1*, conferring DDT resistance, correlated with insertion of a long terminal repeat (LTR) of an Accord retrotransposon in the regulatory region of this gene [24, 25]. However, contrary to other insertion elements linked with over-expression identified so far, this 6.5kb insertion does not contain a transposable element but mainly putative cis-regulatory elements [13].

### 2-Role of 6.5kb and aggravation of pyrethroid resistance

The design a simple PCR diagnostic assay to genotype the 6.5kb SV has allowed to establish its contribution to the resistance phenotype. First, it has been shown that the 6.5kb SV segregates independently from *CYP6P9a* and *CYP6P9b* and thus that it is an additional genetic factor driving pyrethroid resistance beside the allelic variation of both genes previously reported [11]. The independent segregation of the 6.5kb SV is also shown by the increased pyrethroid resistance that it confers either when using WHO bioassays or cone assays. The fixation of this SV besides the resistant alleles of *CYP6P9a* and *CYP6P9b* could explain the resistance escalation currently reported in several mosquito populations of *An. funestus* such as in Mozambique [26] and Malawi [27] and which is reducing the efficacy of insecticide-treated nets [26]. The near fixation of the 6.5kb SV seen here in South Mozambique post bed nets (2016) while it was absent before bed net scale up (2002) is an indication that this SV is strongly associated with resistance exacerbation. The fixation of the 6.5kb in highly resistant wild populations also suggests that escalation of pyrethroid resistance could be driven, among others, by an increased metabolic resistance through genetic elements such as enhancers.

The strong association observed between this structural variation and pyrethroid resistance either with WHO bioassays or cone assays shows that this SV can be used as a resistance marker for monitoring pyrethroid resistance. This comes to add up to the *CYP6P9a* and *CYP6P9b* [14] markers previously identified in the promoters of these genes. This novel assay is even simpler than the PCR-RFLPs previously designed for *CYP6P9a* and *CYP6P9b* as it does not require restriction enzymes.

The detection of this 6.5kb enhancer in the cis-regulatory region of major resistance genes further supports that genetic variations in this region play a major role in driving metabolic resistance as seen for several resistance genes including the P450 *CYP9M10* in *Culex quinquefasciatus* [28] *GSTe2* in *An. funestus* (Mugenzi et al submitted), *CYP6G1* in Drosophila [24]. Therefore, cis-regulatory region of major metabolic resistance genes should be thoroughly investigated to identify more markers to design simple DNA-based assay to detect such resistance in mosquitoes.

### 3. Geographical distribution confirms restriction of gene flow in *An. funestus*

The geographical distribution of the 6.5kb across Africa mirrors closely that of the resistance alleles of both *CYP6P9a* and *CYP6P9b* P450s [13, 14] with the highest frequency observed from southern Mozambique (100%) up to Eastern DRC (72.2%). It also correlates with the Africa-wide distribution of the N485 Ace-1 carbamate resistance allele in this species [29] supporting the existence of an insecticide resistance front in southern Africa which significantly differs to other regions. The South/North clinal increase of frequency in the spread of this SV supports that this resistance likely originated in Southern Mozambique and gradually migrated northwards in combination with local selection forces. The absence of 6.5kb SV in other African regions is in line with barriers of gene flow between populations in this species across the continent with southern African populations consistently differentiated from other regions [30-32]. In contrast, the 6.5kb SV distribution is opposite to that of other markers in this species notably the L119F-GSTe2 conferring DDT/pyrethroid resistance [9] and the A296S-RDL dieldrin resistance marker [33]. Because the 6.5kb SV allele was completely absent in 2002 before LLIN scale up while both *CYP6P9a* and *CYP6P9b* were already detectable although at low frequency (10.9 % and 5.2%) it is likely that this SV was selected later as resistance level increased potential after greater selection pressure. Analysis of larger temporal samples will further help to track the selection of this SV.

Analysis of the distribution of the 6.5kb SV and that of *CYP6P9a* and *CYP6P9b* in natural populations revealed that alleles at these markers do segregate independently in the field as also seen in the hybrid strain FUMOZ-R/FANG at F4 where most of the genotypic combinations were observed. This suggests that regardless of the close proximity of the 3 loci (a total span of 12kb), the three loci are not physically linked. The higher linkage frequency observed in Mozambique and Tanzania can be due to stronger insecticide selection applied against these populations as a consequence of the scale-up of vector control interventions [31]. To determine which gene between *CYP6P9a* and *CYP6P9b* is more linked to the 6.5kb SV, the percentage linkage for *CYP6P9a* and the 6.5kb SV was compared with that for *CYP6P9b* and the 6.5kb SV in the DRC-Mikalayi and Tanzania. For DRC-Mikalayi, *CYP6P9a* and the 6.5kb SV had more identical genotypes (12%) than for *CYP6P9b* and the 6.5kb SV (8%) while in Tanzania, only *CYP6P9a* and the 6.5kb SV identical genotypes (16%) were identified and none for *CYP6P9b* and the 6.5kb SV (0%). Hence this SV although impacting both genes as shown by the comparative qRT-PCR, appears to have a greater linkage to *CYP6P9a*. This could also support a higher fold-change observed for *CYP6P9a* in the field in southern Africa [2, 13, 19]. This could be due to the fact that the 6.5kb is located upstream of the 5’ UTR of *CYP6P9a* but downstream of the 3’UTR of *CYP6P9b*.

### 4. The 6.5kb aggravates the loss of efficacy of insecticide-treated nets

The design of the simple PCR-based assay to genotype the 6.5kb enabled us to assess the impact of such structural variation on the efficacy of insecticide-treated nets including the pyrethroid-only and the PBO-synergist nets. The greater reduction of efficacy that the 6.5kb was shown to cause on pyrethroid-only nets than PBO-based nets is similar to that observed with *CYP6P9a* [13] and *CYP6P9b* [14] in terms of reduction of mortality rate and blood feeding inhibition. It further supports the view that PBO-based nets should be more deployed to control P450-based metabolically resistant mosquito populations such as *An. funestus* throughout southern Africa. The fact that triple homozygote resistant mosquitoes have a greater ability to survive exposure to LLINs shows that resistance escalation in field populations of vectors is likely to significantly reduce the effectiveness of current pyrethroid-based LLINs as recently shown for a population of *An. funestus* in southern Mozambique which was even able to survive exposure to some PBO-based nets [26] due partially to the over-expression of *CYP6P9a* and *CYP6P9b* and the fixation of all three resistance alleles (6.5kb SV, CYP6P9a-R and CYP6P9b-R). The availability of additional DNA-based markers such as these will now enable control programs to assess how resistance is impacting the efficacy of the bed nets in their country and decide whether to adopt PBO-based nets or even new generation nets with another class of insecticide than just pyrethroids.

In conclusion, by elucidating the role of a 6.5kb structural variant in the pyrethroid resistance in *An. funestus*, this study highlighted the important contribution of structural variations in cis-regulatory regions in metabolic resistance in mosquitoes. It also highlighted the role of enhancers in the over-expression of metabolic resistance genes. This study designed a simple molecular diagnostic assay to easily monitor this P450-based metabolic resistance in wild populations. The additive resistance confers by this 6.5kb in the presence of cis-regulatory promoter factors at *CYP6P9a* and *CYP6P9b* as well as allelic variation in coding regions [11] highlights the complex array of evolutionary tools that mosquitoes deploy to survive the scale up of insecticide-based control interventions such as LLINs. The near fixation of triple resistant (SV/CYP6P9a/CYP6P9b) in southern Africa calls for urgent action in using alternative insecticides than pyrethroids for IRS and new generation LLINs with PBO and more preferably with new another insecticide class.

## Materials and methods

### Mosquito laboratory strains and field populations

Two *An. funestus* laboratory colonies of FUMOZ-R and FANG, which are resistant and susceptible to pyrethroids respectively, were used [34]. Available DNA samples from previous studies including F_8_ generation samples derived from the reciprocal crosses between the FUMOZ and FANG [17] were used. Additionally, field collected mosquito samples from several countries across Africa were also used for genotyping including Democratic Republic of Congo (DRC) (2015), Cameroon (2016) for central Africa, Ghana (2014) for West Africa, Uganda (2014) and Tanzania (2015) for East Africa, Zambia (2015), Malawi (2014) and Mozambique (2016) for Southern Africa. These field samples were collected indoor using electric aspirators as previously described [13].

Furthermore, reciprocal crosses between FUMOZ-R and FANG were carried out to assess the correlation between the 6.5kb SV and pyrethroid resistance phenotype. To perform these crosses, 30 males FUMOZ-R and 30 virgin females FANG where allowed to mate and the eggs reared to the next generation and adults of following generations inter-crossed until up to the F_6_ generation.

### Insecticide susceptibility assays

The resistance level of mosquitoes from F_4_ to F_6_ generation was tested using WHO bioassays performed according to WHO protocols [35]. Briefly, 2-5 day-old, unfed female mosquitoes were exposed for 1h to papers impregnated with the pyrethroids permethrin (0.75%) and deltamethrin (0.05%). Moreover, these mosquitoes were also exposed to the same papers for 30 min and 90 min to generate highly susceptible and highly resistant individuals. An untreated paper was also used as negative control.

### WHO cone bioassays

The efficacy of insecticide-treated net against the strains used was tested using cone assay following the WHO protocol [36]. Mosquitoes from the 6^th^ generation of the FUMOZ-R and FANG crosses were tested against two WHO recommended LLINs PermaNet 2.0 and PermaNet 3.0. PermNet 2.0 nets consist of polyester coated with the deltamethrin to a target dose of 55 mg/m^2^ (±25%) while PermaNet 3.0 has a higher dose of deltamethrin of 85 mg/m^2^ (±25%) and piperonyl butoxide (PBO) a synergist incorporated with deltamethrin in the polyethylene roof (Vestergaard, Frandsen, Lausanne, Switzerland). Briefly, 5 replicates of 10 F_6_ females (2–5 days old) were placed in plastic cones attached to 30cm x 30cm pieces of nets PermaNet 2.0, PermaNet 3.0 (side) and PermaNet 3.0 (roof), and an insecticide free net as a control. After 3 minutes exposure, the mosquitoes were transferred in holding paper cups and provided with cotton soaked in 10% sugar solution. Mortality was recorded 24 hours post exposure.

### Confirmation of the presence of 6.5kb SV from whole genome sequences from different regions of Africa

The Pool-Seq data from several populations of *An. funestus* were analysed to confirm the presence and distribution of the 6.5kb insert Africa-wide following protocol described recently [13]. Initial processing and quality assessment of the sequence data was performed as described recently [13]. Pool-Seq R1/R2 read pairs and R0 reads were aligned to the reference sequence using bowtie2 [37]. Variant calling was carried out using SNVer version 0.5.3, with default parameters [38]. SNPs were filtered to remove those with total coverage depth less than 10 and more the 95^th^ centile for each sample as the allele frequency estimates could be inaccurate due to low coverage or misaligned paralogous sequence, respectively.

### Amplification and cloning of *CYP6P9a* and *CYP6P9b* intergenic region

The Livak protocol [39] was used to extract DNA from the collected samples. The extracted DNA was quantified using NanoDrop lite™ spectrophotometer (Thermo Scientifc, Wilmington, USA). The intergenic region between *CYP6P9a* and *CYP6P9b* was amplified for both the pyrethroid resistant lab strain FUMOZ-R and the FANG pyrethroid susceptible lab strain. The intergenic region for the FANG was amplified using 6P9a5F and GAP3R (S1) primers using the 1U KAPA Taq polymerase (Kapa Biosystems, Boston, MA, USA) in 1X buffer A, 25mM MgCl_2_, 25mM dNTPs, 10 mM of each primer was used to constitute a 15 µL PCR reaction mix using the following conditions, initial denaturation step of 3 min at 95 °C, followed by 35 cycles of 30 s at 94 °C, 2 minutes s at 58 °C, and 60 s at 72 °C, with 10 minutes at 72 °C for final extension. The FUMOZ-R was amplified with GAP1F and GAP3R (S1) primers using Phusion high-fidelity DNA polymerase (Thermos Scientific, Waltham, Massachusetts, United States). The mix was made using the GC buffer, 3% DMSO, 25mM dNTPs, 10 mM of each primer and DNA from FUMOZ as template. The PCR conditions were as follows 10 minutes at 98°C pre-denaturation, 35 cycles 10 seconds at 98°C denaturation, 30 seconds at 62°C annealing, 4 minutes at 72°C extension and a final extension at 72°C for 10 minutes. The PCR product was run on a 1% gel and visualized on ENDURO™GDS (labnet, Edison, New Jersey, USA) UV transilluminator. The PCR product was gel purified and ligated to PJET1.2 blunt end vector and sequenced. Sequencing data were analysed with Bioedit [40].

### Cloning of the intergenic *CYP6P9a-CYP6P9b for* promoter activity assay

The 8.2 kb intergenic region of *CYP6P9a* and *CYP6P9b* of the FUMOZ-R colony was amplified using the primers 6P9bUTR8.2F CGA GCT CGT AAG TAA CAC ACA AAA TGG T and 6P9aUTR8.2R CGG CTA GCC GAT TTC GTT CGC CGA ATT CCA, using Phusion high-fidelity DNA polymerase (Thermo Scientific) with the GC buffer, 3% DMSO, 25mM dNTPs, 10 mM of each primer and DNA from FUMOZ-R as template. The PCR conditions were as follows; 10 minutes at 98°C pre-denaturation, 35 cycles 10 seconds at 98°C denaturation, 30 seconds at 62°C annealing, 4 minutes at 72°C extension and a final extension at 72°C for 10 minutes. The PCR product was run on a 1% gel and visualized on UV transilluminator. The band at 8.2 kb was gel purified and ligated to pjet1.2 vector (Figure S3a). The recombinant plasmid was digested with Sac1 and Nde1 restriction enzyme and sub-cloned in pGL3 Basic luciferase vector (Promega, Madison, USA).

### Luciferase reporter assay

The transfection and the luciferase assay was done as previously described [13] using *An. gambiae* 4a-2 cell line (MRA-917 MR4, ATCC® Manassas Virginia). Due to the large difference in size between the recombinant promoter constructs, equimolar amount of each constructs was used for the transfection. Luciferase assay compared the 8.2 kb intergenic of the FUMOZ-R to the 0.8 kb *CYP6P9a* promoter constructs for FUMOZ-R and FANG. Five independent transfections were carried out. The 0.8kb *CYP6P9a* recombinant promoter construct of FUMOZ-R contained the core promoter of *CYP6P9a* before the insertion point. Briefly, 600 ng of each promoter construct were transfected using the effectene transfection reagent (Qiagen, Hilden, Germany). Two additional plasmids were used, the LRIM promoter construct which served as a positive control and the actin-Renilla internal control which is used for normalization. A luminometer (EG & G Berthold, Wildbad, Germany) was used to measure luciferase activity two days post transfection.

### Development of a PCR for genotyping the 6.5 kb structural variant between *CYP6P9a* and *CYP6P9b*

A PCR was designed to discriminate between mosquitoes with the 8.2kb (FUMOZ-R) and 1.7kb (FANG*) CYP6P9a* and *CYP6P9b* intergenic region. Briefly three primers where designed, two (FG_5F: CTT CAC GTC AAA GTC CGT AT and FG_3R: TTT CGG AAA ACA TCC TCA A) at region flanking the insertion point and a third primer (FZ_INS5R: ATATGCCACGAAGGAAGCAG) in the 6.5kb insertion (Figure 2b). 1U KAPA Taq polymerase (Kapa Biosystems, Boston, MA, USA) in 1X buffer A, 25mM MgCl_2_, 25mM dNTPs, 10 mM of each primer was used to constitute a 15 µL PCR reaction mix using the following conditions; initial denaturation step of 3 min at 95 °C, followed by 35 cycles of 30 s at 94 °C, 30 s at 58 °C, and 60 s at 72 °C, with 10 min at 72 °C for final extension. The amplicon was revolved on a 1.5% agarose gel stained with Midori Green Advance DNA Stain (Nippon genetics Europe GmbH) and revealed on UV transilluminator.

To assess the ability of the assay to discriminate between mosquitoes with the structural variant and those without, the assay was tried on genomic DNA extracted from 48 FUMOZ-R and 50 FANG female mosquitoes. And later on, mosquitoes for 8^th^ generation of crosses between FUMOZ-R and FANG which had been exposed to 0.75% permethrin for 180 min (alive providing the highly resistant) and 30 min (the dead being the highly susceptible) were genotyped for this SV to assess the correlation between the insertion genotype and the insecticide susceptibility phenotype.

### Correlation between the 6.5kb PCR assay and the *CYP6P9a/b* PCR-RFLP

The efficacy of the newly designed PCR for the 6.5kb SV was compared with the PCR-RFLP methods for detecting *CYP6P9a* and *CYP6P9b* resistant allele to check the association between the 6.5k SV and each of the P450s and also for the combined effect of these three markers as described in previously [13, 14]. DNA samples from previous studies were used as test materials to compare the three assays.

### Quantitative real time PCR

Total RNA was extracted from F_2_ generation of FUMOZ-R and FANG crosses which had been genotyped using the newly designed 6.5k SV detection PCR and grouped in three sets: homozygous for 6.5kb SV (SV+/SV+), heterozygous (SV+/SV-) and homozygous without the SV (SV-/SV-). DNA was extracted from the legs. Briefly 4 to 6 legs were pulled from each mosquito and placed in 1.5 ml tube. 25μl of 1X PCR buffer B (Kapa Biosystems, Boston, MA, USA) pre-warmed at 65°C was added then the legs were ground. After the grinding the tubes were centrifuged at 13000rpm for 2 minutes. The samples were then incubated at 95°C for 30 minutes in a thermocycler. The remaining body of the mosquito was kept in RNAlater individually. From each group, 30 mosquitoes were pooled in sets of 10 females and RNA was extracted using the Arcturus PicoPure RNA Isolation Kit (Life Technologies Carlsbad, California, United States). cDNA were synthesised from each RNA pool using the Superscript III (Invitrogen, Carlsbad, California, United States) as previously described [27]. The expression pattern of *CYP6P9a* and *CYP6P9b* was assessed by quantitative real time PCR (qRT-PCR) (Agilent MX3005) to assess the correlation between the presence of the 6.5kb SV and the expression pattern of these two resistance genes.

### Experimental hut trials

To assess the association between the 6.5kb SV and a potential reduction in the efficacy of insecticide treated nets, an experimental hut trial was performed in a field station (possessing 12 huts) located at Mibellon (6°4′60′′N, 11°30′0′′E), a village in Cameroon located in the Adamawa Region; Mayo Banyo division and Bankim Sub-division. A release recapture experiment was done using 5^th^ and 6^th^ generations of the FANG/FUMOZ-R crosses where about 50 to100 female mosquitoes were released in each hut in the evening at 8:00pm with a volunteer sleeping under each net. Both alive and dead, blood fed and unfed mosquitoes were collected in the morning.

### Statistical analysis

Statistical analysis including student’s t test, Fisher exact test and odd ratio were done using the online MEDCALC (https://www.medcalc.org/index.php), vassarstats (http://vassarstats.net/) and GraphPad prism 7.05 softwares

## Funding

This work was supported by Wellcome Trust Senior Research Fellowship in Biomedical Sciences to Charles S. Wondji (101893/Z/13/Z and 217188/Z/19/Z).

## Acknowledgments

The authors will like to thank the inhabitants of Mibellon for their support during the study.

## Conflicts of Interest

The authors declare that they have no competing interests.

---

**S1 Fig:**
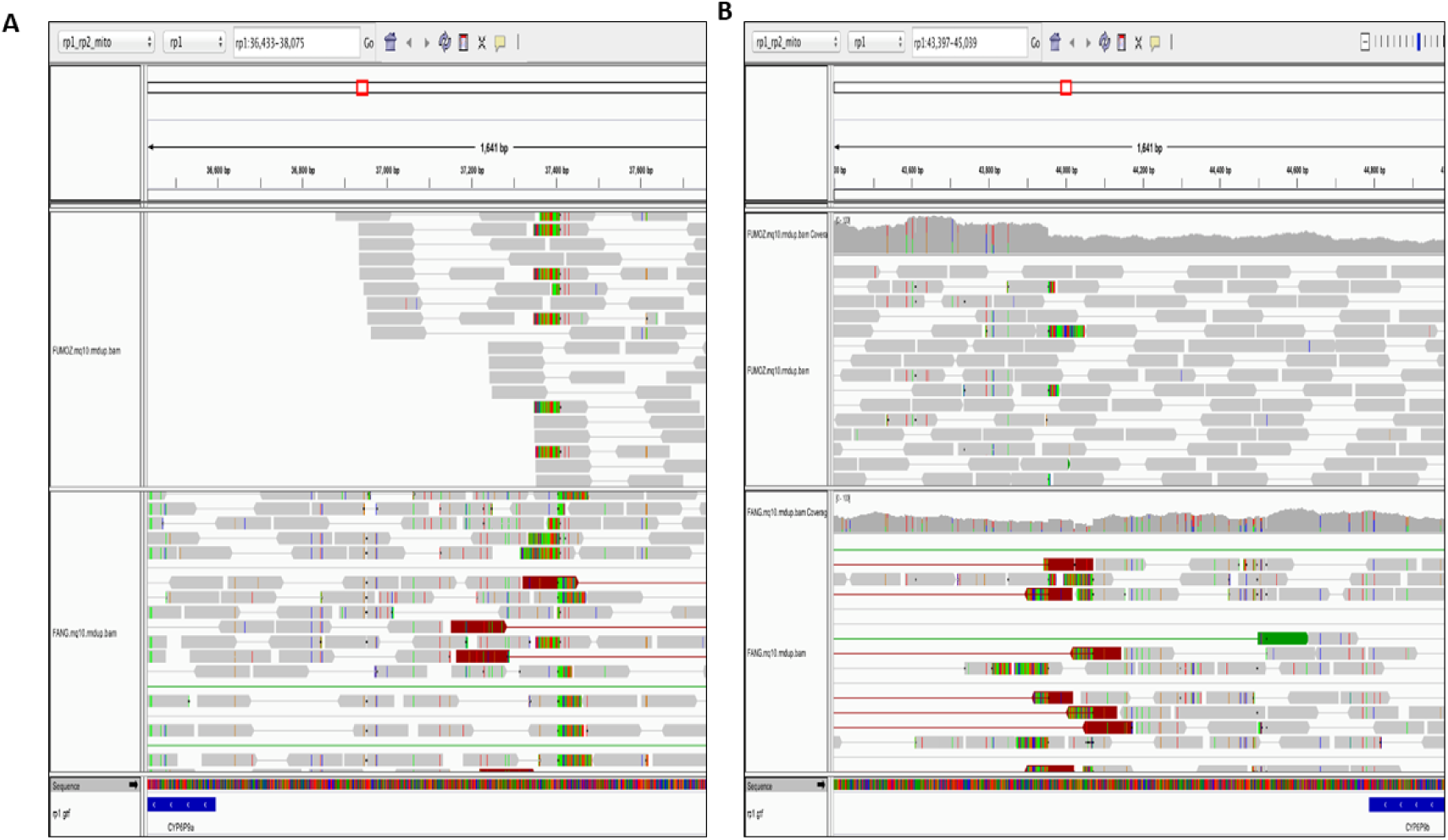
Insertion of a 6.5kb intergenic fragment between *CYP6P9a* and *CYP6P9b* in southern African mosquitoes. (A) IGV screenshot showing the left breakpoint of the insertion in FUMOZ-R (upper) and FANG (lower) reads. The large number of “soft-clipped” reads (all clipped bases shown in colour in the reads) indicates the putative breakpoint. In FUMOZ, some reads are left-clipped at BAC sequence positions 37409/37410. In FANG, some reads are left-clipped at the same position, and some right-clipped at 37404/37405. (B) IGV screenshot showing the right breakpoint of the insertion in FUMOZ-R (upper) and FANG (lower) reads. In FUMOZ, some reads are right-clipped at BAC positions 43954/43955. In FANG, some reads are right-clipped and some left-clipped at the same positions. In addition, in FANG there are right-clipped reads at 44053/44054 and left-clipped reads at 44070/44071.

**S2 Fig:**
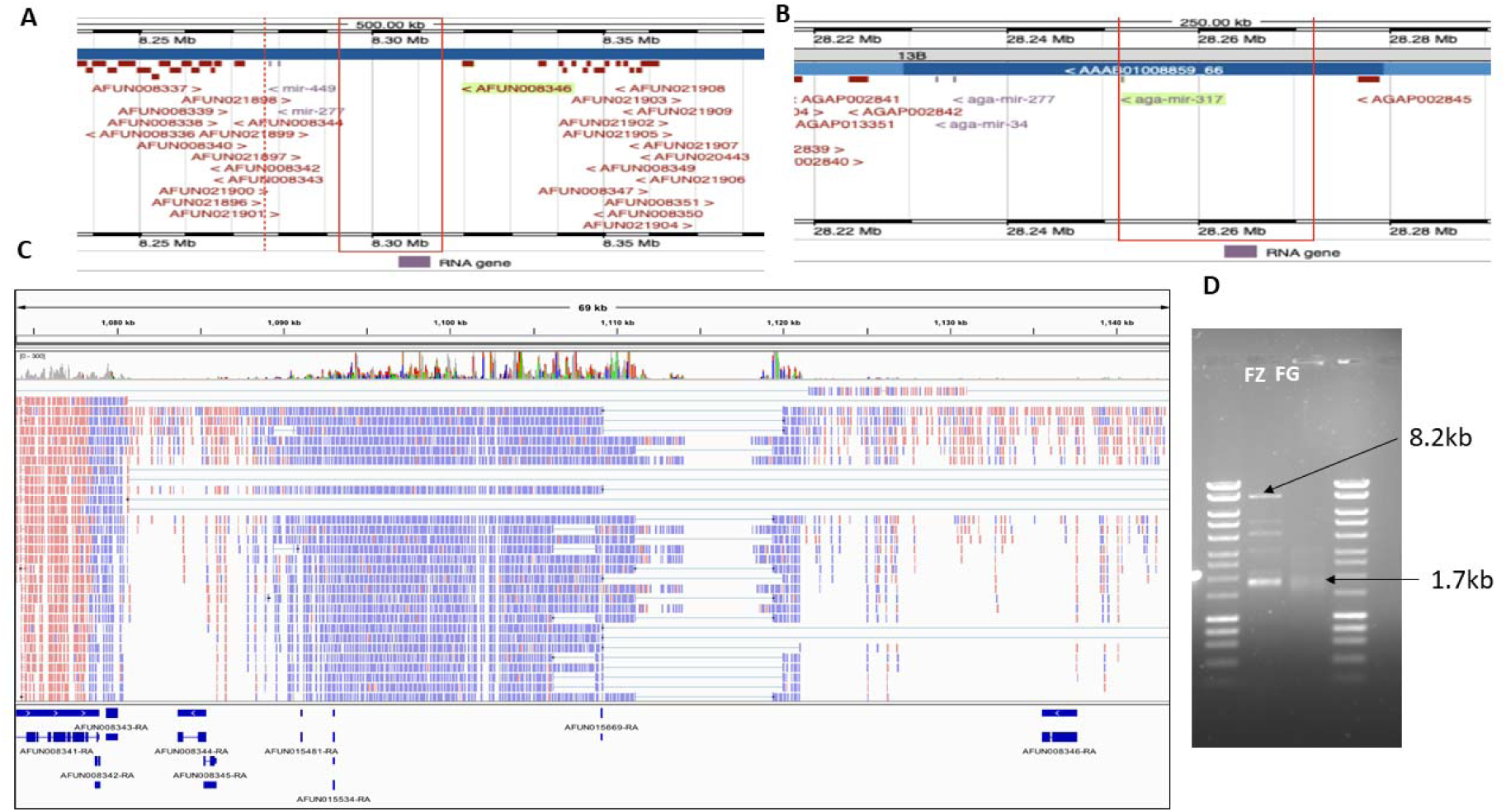
Genomic context of the “parent” region from which the CYP6 insertion was derived. (A) *Anopheles funestus* CM012071.1 :8,296,288-8,555,956 at the top, showing the region inserted into the CYP6 cluster in the red box. (B) is the orthologous region in *Anopheles gambiae* at chromosome 2R:28217050-28291000. (C) The 6.5 kb insertion comes from a transcribed region of the genome. IGV screenshot showing a representative RNAseq alignment (FUMOZ) against the AfunF1 genome assembly in the insertion “parent” region. Annotated genes are indicated by blue boxes at the bottom of the image. RNAseq reads aligned to the negative strand are shown in blue and to the positive strand in red. Thin lines indicate reads spanning an intron. A large region, including three micro-RNAs, is transcribed on the negative strand. Splicing is seen near the 5’ end, around mir-317. The true pattern of splicing may be hidden by the assembly gap within the transcribed region. (D) PCR amplification of the 8.2kb (FUMOZ-R) and 1.7kb (FANG) intergenic fragment between CYP6P9a and CYP6P9b for subsequent cloning and sequencing.

**S3 Fig:**
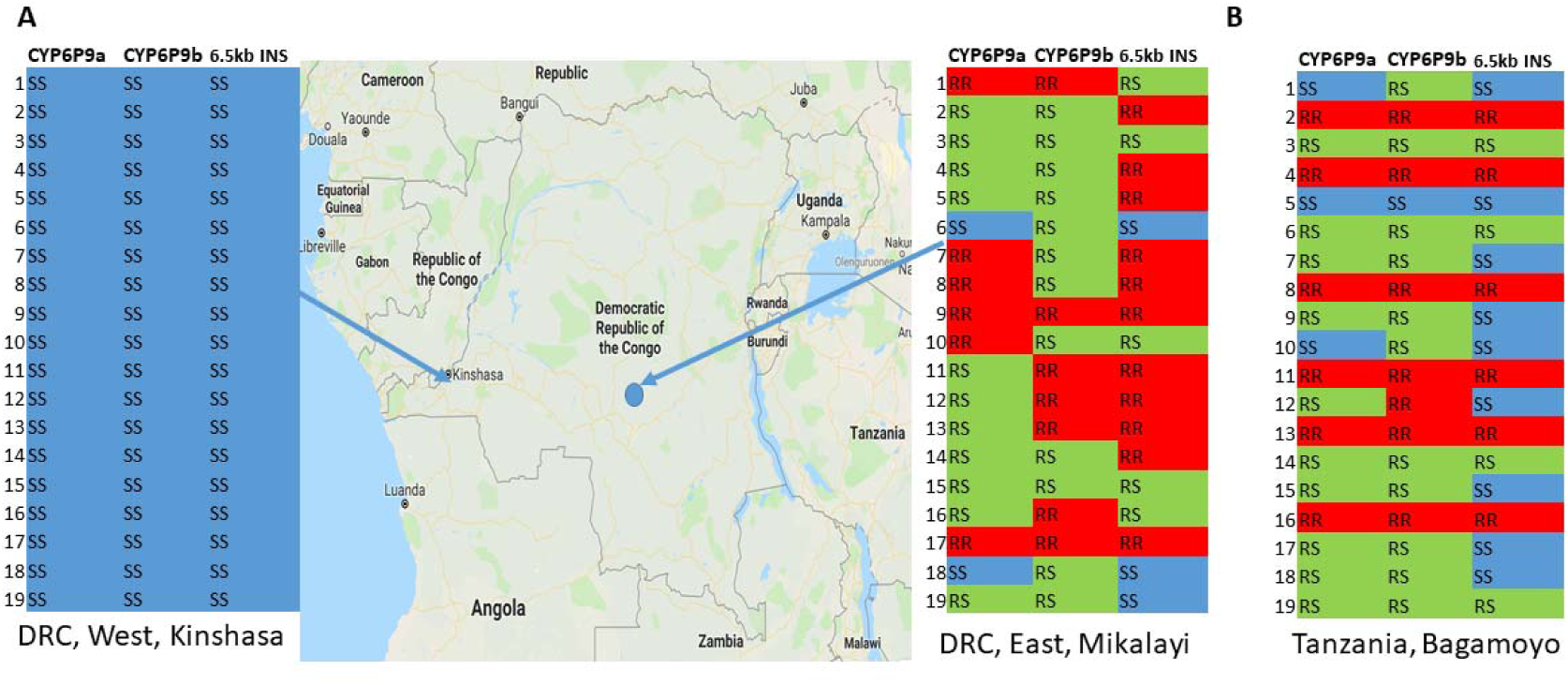
Illustration of the segregation of the genotypes of the 6.5kb SV with those of the duplicated P450 *CYP6P9a* and *CYP6P9b* in Democratic Republic of Congo (DRC) and Tanzania. (A) Contrasting genotypic distribution between West (Kinshasa) and East (Mikalayi) populations from DRC. (B) Segregation of genotypes at the three loci in Bagamoyo in Tanzania.

**S4 Fig:**
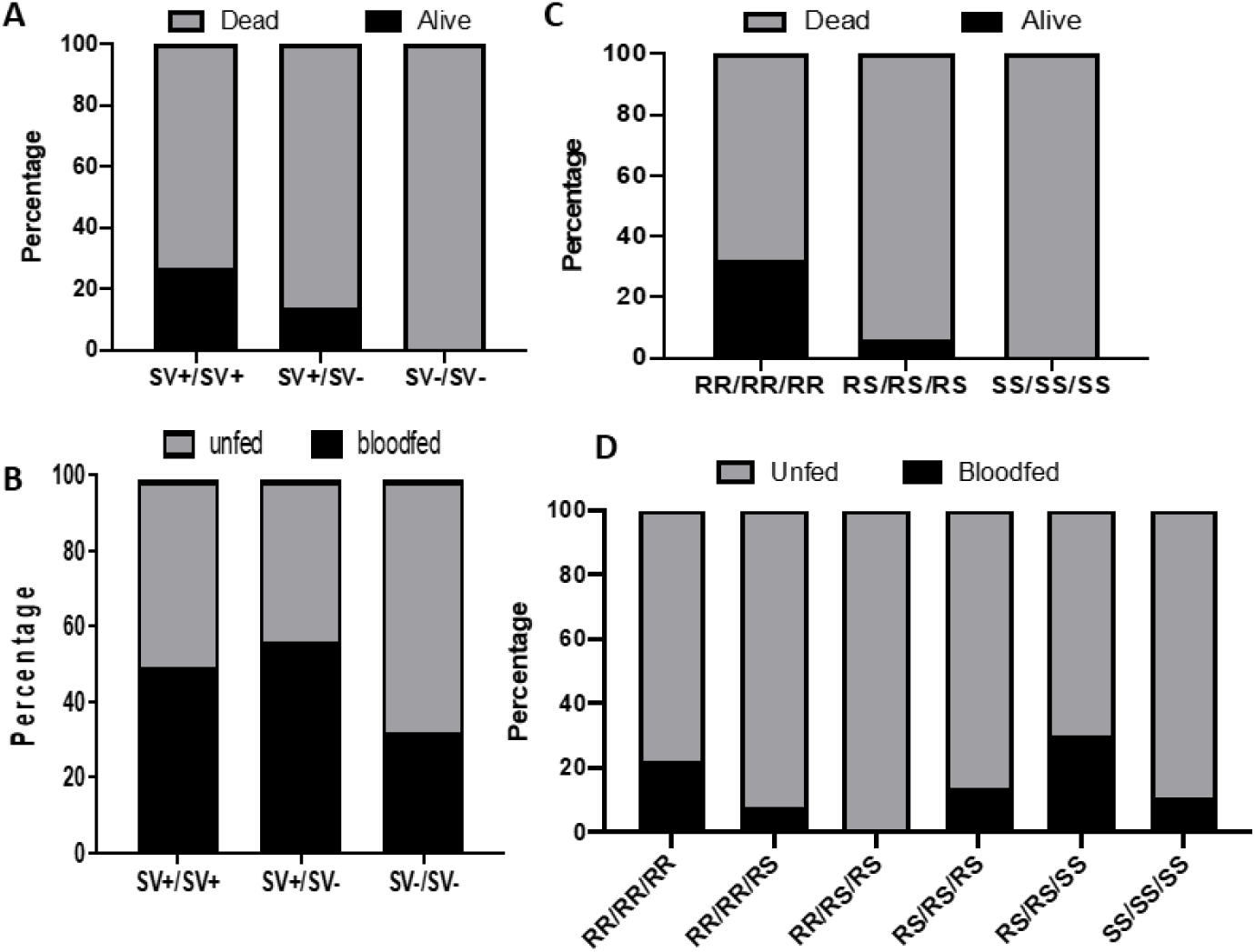
Impact of the 6.5kb enhancer on the ability to survive and blood feed in the presence of PermaNet 2.0 and PermaNet 3.0 in experimental hut trials: (A) Distribution of the 6.5kb SV among mosquitoes alive and dead after exposure to PermaNet 3.0 (B) Distribution of the 6.5kb *SV* genotypes between mosquitoes that bloodfed in presence of PermaNet 2.0. (C) Distribution of the combined genotype for the *CYP6P9a, CYP6P9b* and 6.5kb SV among mosquitoes alive and dead after exposure to PermaNet 3.0. (D) Prevalence of combined genotypes of *CYP6P9a, CYP6P9b* and 6.5kb SV in blood-fed and unfed mosquitoes for PermaNet 2.0 showing no significant difference.

**S5 Fig:**
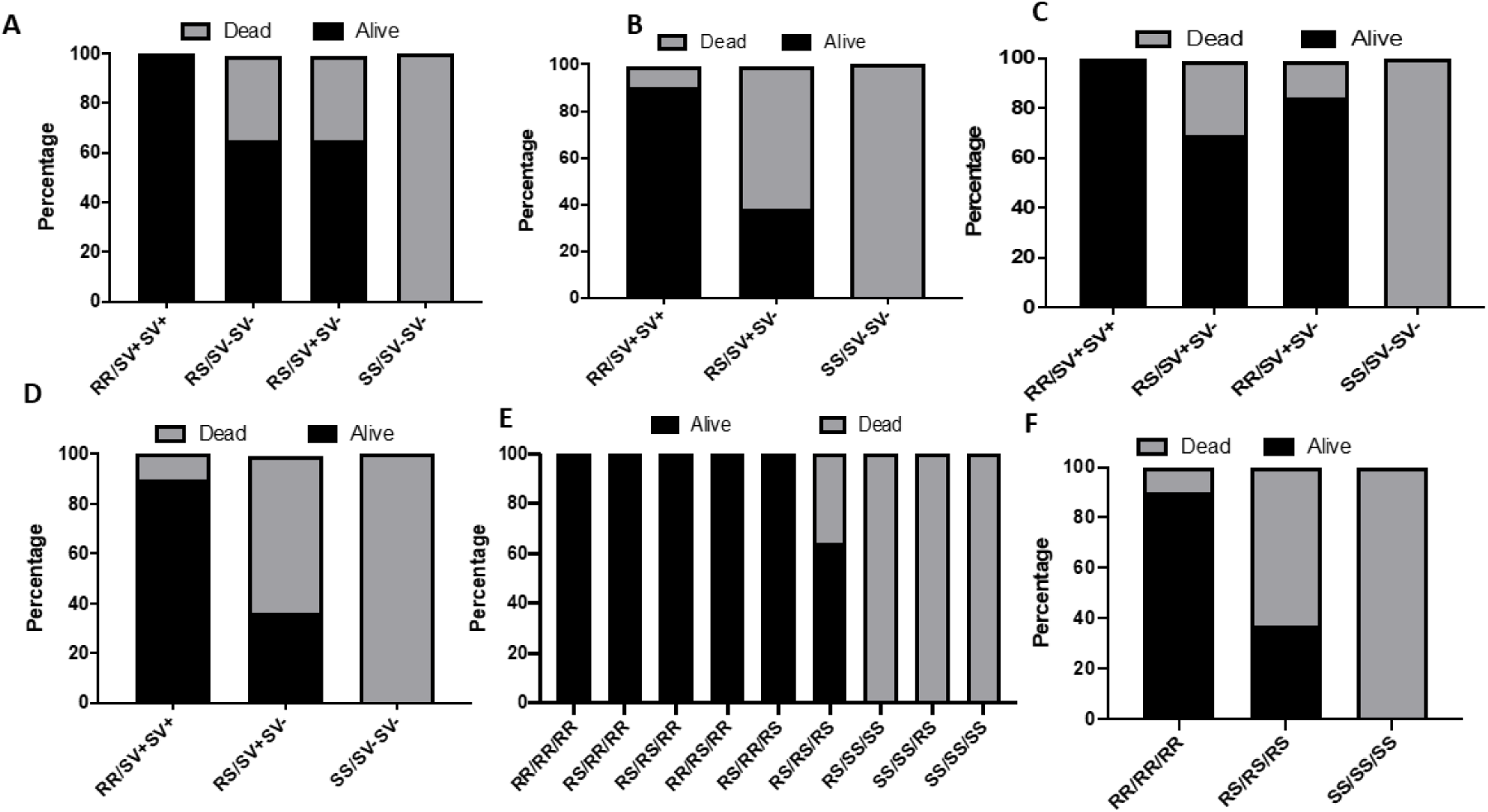
Combined effects of 6.5kb and each of the P450 genes *CYP6P9a* and *CYP6P9b* using cone assays: (A) Distribution of the combined genotypes of 6.5kb SV and *CYP6P9a* showing that both genotypes combined to increase the ability to withstand PermaNet 2.0 exposure and (B) for PermaNet 3.0. (C) is the same for SV with *CYP6P9b* for PermaNet 2.0 also showing that both SV/CYP6P9b genotypes combined to increase the ability to withstand PermaNet 2.0 exposure. (D) Distribution of the combined genotypes of 6.5kb SV and *CYP6P9b* showing that both genotypes combined to increase the ability to withstand exposure to PermaNet 3.0 (side). (E) Distribution of the combined genotypes for the *CYP6P9a, CYP6P9b* and 6.5kb SV among mosquitoes alive and dead after exposure to PermaNet 2.0. (F) is the distribution for the combined genotypes of all three loci for PermaNet 3.0 (side).

**S6 Fig:**
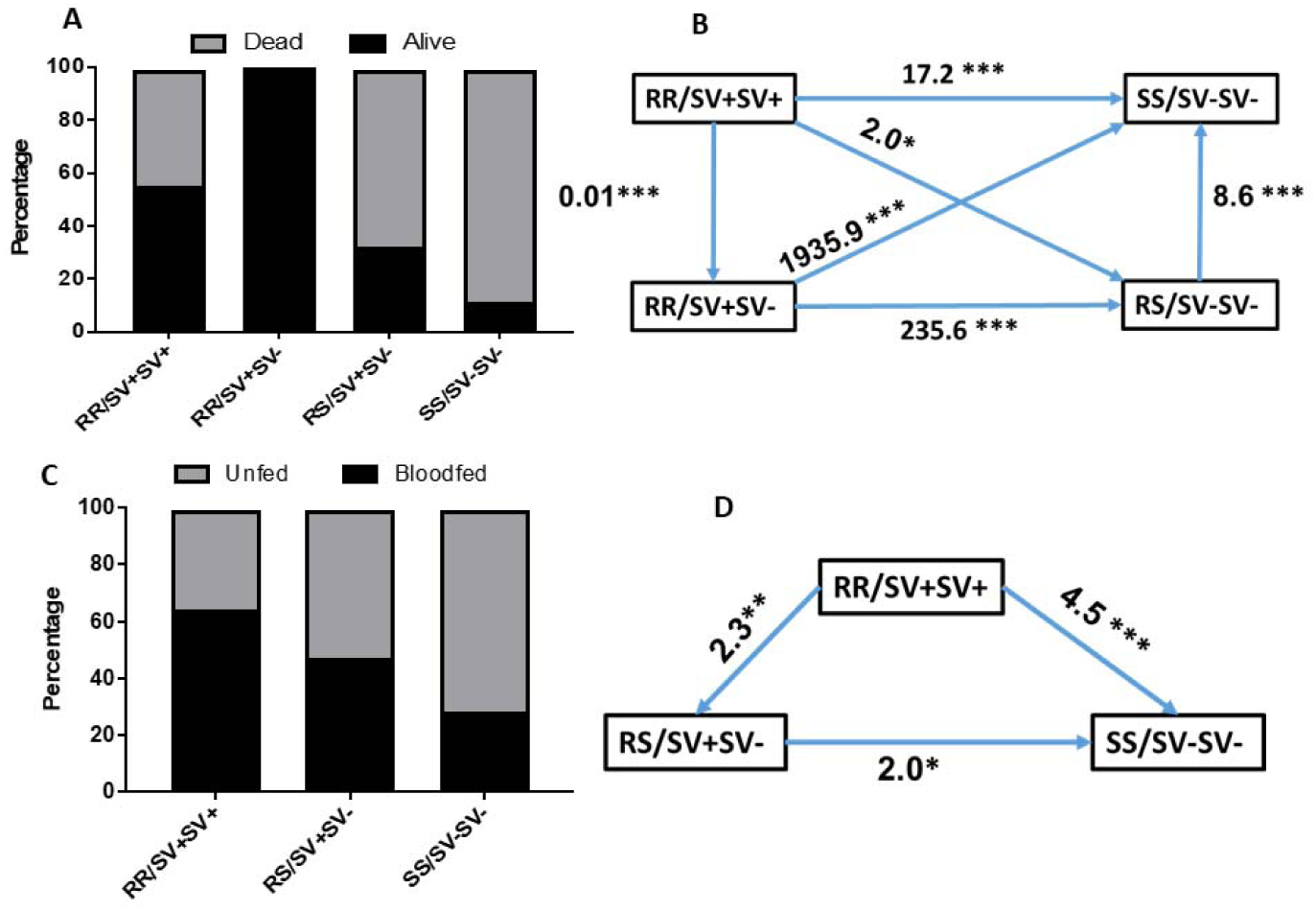
*6.5kb SV* combines with *CYP6P9a* to further reduce the efficacy of insecticide-treated nets. (A) Distribution of the combined genotypes of both *CYP6P9a* and 6.5kb SV showing that genotypes at both genotypes combined to additively increase the ability to survive after exposure to PermaNet 2.0. (B) Ability to survive exposure to PermaNet 2.0 (Odds ratio) of the double homozygote resistant (RR/SV+SV+) genotypes of *CYP6P9a* and 6.5kb *SV* compared to other genotypes supporting the additive resistance effect of both genes. (C) Distribution of the combined genotypes of both *CYP6P9a* and *6.5 kb SV* after exposure to PermaNet 3.0 revealing an additive effect of both genes in increasing ability to blood feed. (D) Comparison of blood feeding ability of combined genotypes of *CYP6P9a* and 6.5 kb *SV* showing a significantly higher ability (odd ratio) to blood feed for mosquitoes double homozygote resistant (RR/RR).

**S7 Fig:**
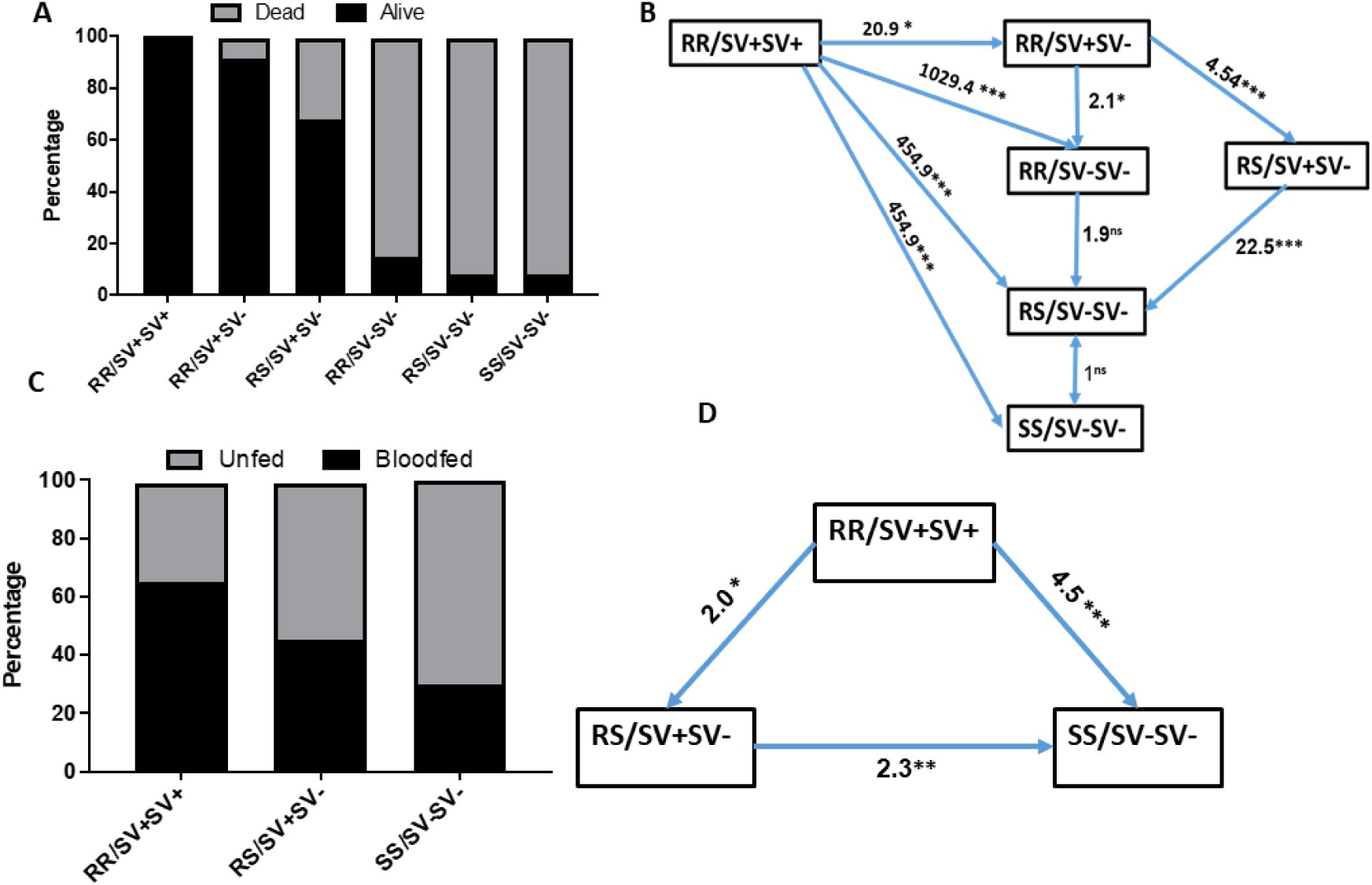
The 6.5kb SV combines with *CYP6P9b* to further reduce the efficacy of insecticide-treated nets. (A) Distribution of the combined genotypes of both *CYP6P9b* and *6.5 kb SV* showing that genotypes at both genotypes combined to additively increase the ability to survive after exposure to PermaNet 2.0. (B) Ability to survive exposure to PermaNet 2.0 (Odds ratio) of the double homozygote resistant (RR/RR) genotypes of *CYP6P9b* and *6.5 kb SV* compared to other genotypes supporting the additive resistance effect of both genes. (C) Distribution of the combined genotypes of both *CYP6P9b* and *6.5 kb SV* after exposure to PermaNet 3.0 revealing an additive effect of both genes in increasing ability to blood feed. (D) Comparison of blood feeding ability of combined genotypes of *CYP6P9b* and *6.5 kb SV* showing a significantly higher ability (odd ratio) to blood feed for mosquitoes double homozygote resistant (RR/RR).

**S8 Fig:**
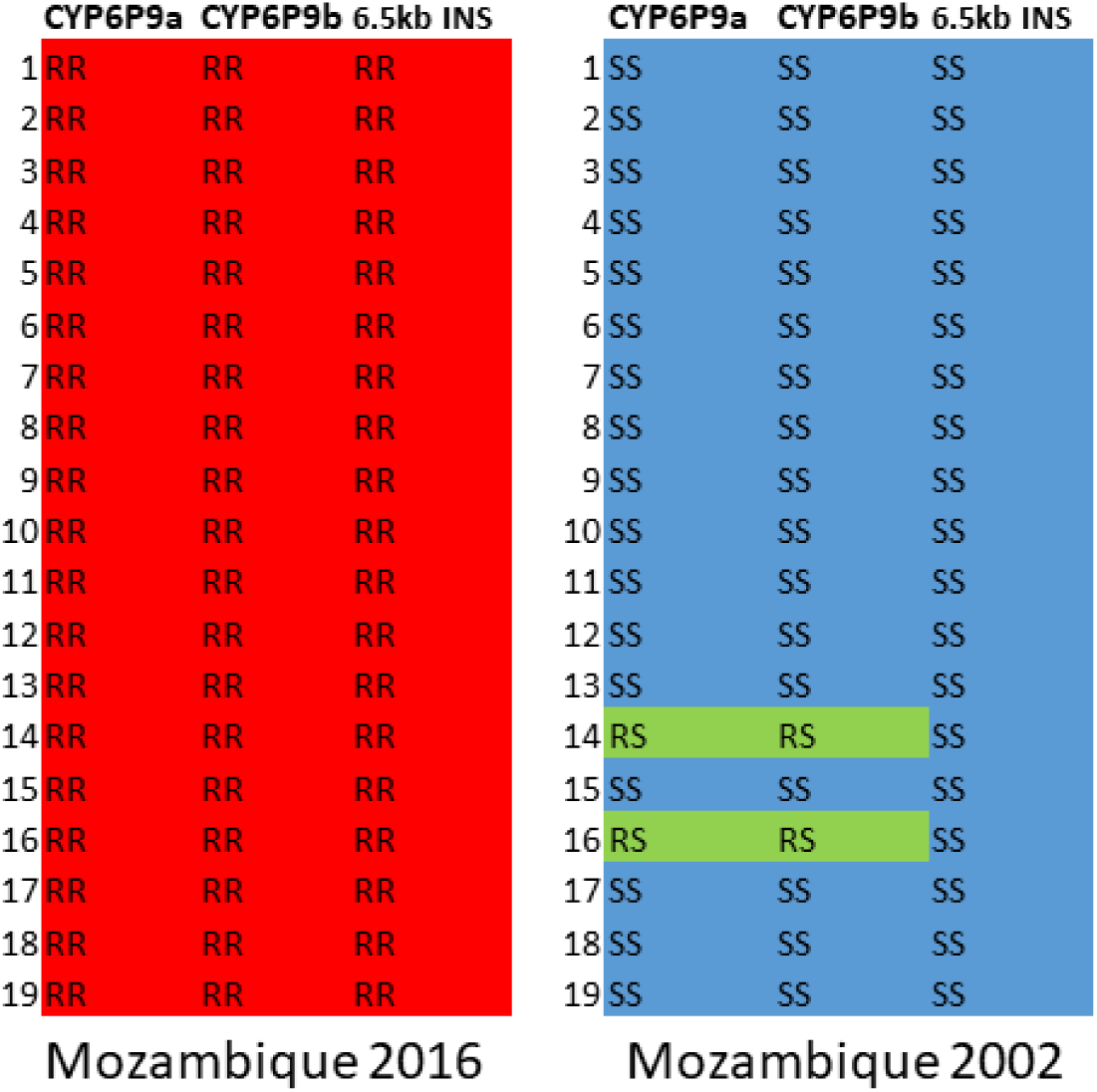
Contrasting distribution of genotypes of the 6.5kb SV and that of CYP6P9a and CYP6P9b before the scale up of bed nets in Mozambique (2002) and after the scale up (2016) showing that these alleles have been selected to fixation in south Mozambique.

**S1 Table:**
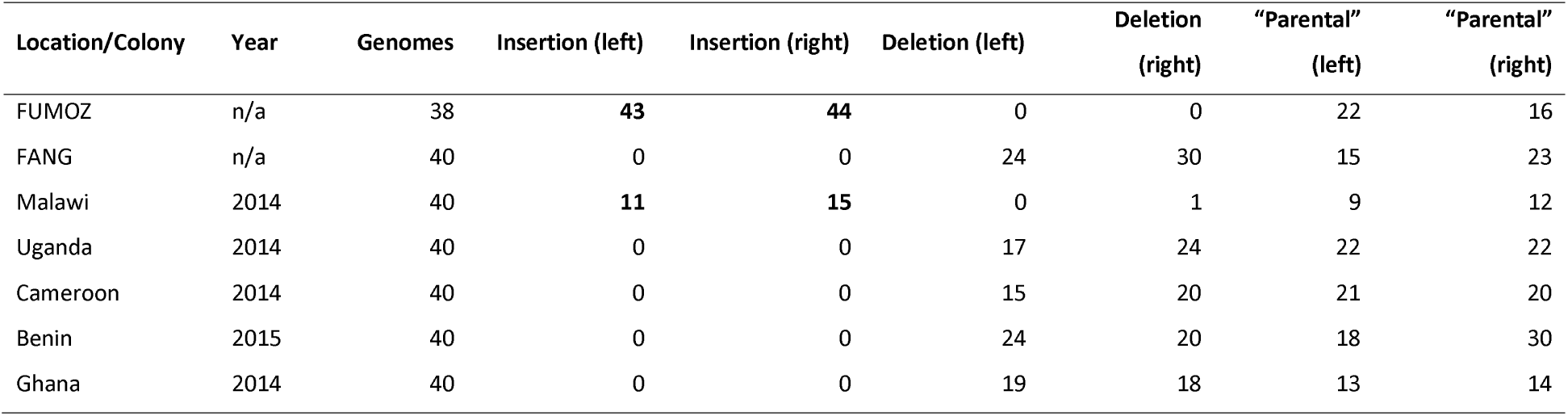
Counts of reads aligned at the left and right breakpoints of the 6.5 kb insertion supporting different haplotypes: “Insertion”=insertion present between *CYP6P9a* and *b*; “Deletion”=insertion absent between *CYP6P9a* and *b*; “Parental”=read originating from elsewhere in the genome, from where the inserted sequence was derived.

**S2 Table:** Correlation between genotypes of the 6.5kb SV and pyrethroid resistance phenotype after WHO bioassays and additive effect to *CYP6P9a* and *CYP6P9b*

| Genotype comparison | OR | CI | P value |
| --- | --- | --- | --- |
| **6.5 kb SV** |  |  |  |
| SV+SV+ vs SV-SV- | 2079.4 | 109.7 -39422.6 | P<0.0001 |
| SV+SV+ vs SV+SV- | 4.8 | 0.2-102.6 | P<0.0001 |
| SV+SV- vs SV-SV- | 600.3 | 106.2-3391.6 | P<0.0001 |
| SV+ vs SV- | 242.4 | 32.3-1821.1 | P<0.0001 |
| **CYP6P9a/SV** |  |  |  |
| RR/SV+SV+ vs SS/SV-SV- | 1300.2 | 59.7-28314.5 | P<0.0001 |
| RS/SV+SV- vs SS/SV-SV- | 1445.5 | 198.3-10536.9 | P<0.0001 |
| RR/SV+SV+ vs RS/SV+SV- | 1.3 | 0.1-30.3 | P=0.8355 |
| **CYP6P9b/SV** |  |  |  |
| RR/SV+SV+ vs SS/SV-SV- | 906.2 | 40.9-20060.3 | P < 0.0001 |
| RS/SV+SV- vs SS/SV-SV- | 4137 | 195.0-87777.5 | P < 0.0001 |
| RR/SV+SV+ vs RS/SV+SV- | 0.219 | 0.0041 to 11.6247 | P = 0.4536 |
| **SV/CYP6P9a/CYP6P9b** |  |  |  |
| RR/RR/SV+SV+ vs SS/SS/SV-SV- | 748.6 | 33.4360 to 16760.4431 | P < 0.0001 |
| RS/RS/SV+SV-RS vs SS/SS/SV-SV- | 3979.4 | 187.5 to 84473.7 | P < 0.0001 |
| RR/RR/SV+SV+ vs RS/RS/SV+SV- | 0.2 | 0.0035 to 10.1 | P = 0.4107 |

**S3 Table:** Correlation between genotypes of the 6.5kb SV and pyrethroid resistance phenotype after PermaNet 2.0 cone assays and additive effect to *CYP6P9a* and *CYP6P9b*

| Genotype comparison | OR | CI | P |
| --- | --- | --- | --- |
| **6.5 kb SV** |  |  |  |
| SV+SV+ vs SV-SV- | 1798.3 | 97.6 -33141.8 | P<0.0001 |
| SV+SV+ vs SV+SV- | 7.4 | 2.6-21.5 | P<0.0002 |
| SV+SV- vs SV-SV- | 261.5 | 15.4-4447.3 | P<0.0001 |
| SV+ vs SV- | 27.7 | 13.0-59.0 | P<0.0001 |
| **SV/CYP6P9a** |  |  |  |
| RR/SV+SV+ vs SS/SV-SV- | 12727 | 248.1-652982.0 | P < 0.0001 |
| RS/SV+SV- vs SS/SV-SV- | 281.9 | 16.5-4827.0 | P = 0.0001 |
| RR/SV+SV+ vs RS/SV+SV- | 45.1 | 2.6-777.4 | P = 0.0087 |
| **SV/CYP6P9b** |  |  |  |
| RR/SV+SV+ vs SS/SV-SV- | 11929 | 232.3-612667.4 | P < 0.0001 |
| RS/SV+SV- vs SS/SV-SV- | 336.1 | 19.5-5780.6 | P = 0.0001 |
| RR/SV+SV+ vs RS/SV+SV- | 35.5 | 2.0-615.4 | P = 0.0142 |
| **SV/CYP6P9a/CYP6P9b** |  |  |  |
| RR/RR/SV+SV+ vs SS/SS/SV-SV- | 10439 | 203.0-536880.0 | P < 0.0001 |
| RS/RS/SV+SV- vs SS/SS/SV-SV- | 243.5 | 14.2-4168.1 | P = 0.0001 |
| RR/RR/SV+SV+ vs RS/RS/SV+SV- | 42.9 | 2.5-740.6 | P = 0.0097 |

**S4 Table:** Correlation between genotypes of the 6.5kb SV and pyrethroid resistance phenotype after PermaNet 3.0 side cone assays and additive effect to *CYP6P9a* and *CYP6P9b*

| Genotype comparison | OR | CI | P |
| --- | --- | --- | --- |
| **6.5 kb SV** |  |  |  |
| SV+SV+ vs SV-SV- | 413.4 | 23.8-7190.2 | <0.0001 |
| SV+SV+ vs SV+SV- | 17.6 | 8.0-38.6 | <0.0001 |
| SV+SV- vs SV-SV- | 24.7 | 1.4-422.0 | 0.0 |
| SV+ vs SV- | 12.1 | 5.8-25.5 | <0.0001 |
| **SV/CYP6P9a** |  |  |  |
| RR/SV+SV+ vs SS/SV-SV- | 374.7 | 21.6-6490.1 | <0.0001 |
| RS/SV+SV- vs SS/SV-SV- | 25.9 | 1.5 to 443.2 | 0.0248 |
| RR/SV+SV+ vs RS/SV+SV- | 15.2 | 6.9 to 33.5 | <0.0001 |
| **SV/CYP6P9b** |  |  |  |
| RR/SV+SV+ vs SS/SV-SV- | 379.7 | 22.0-6575.3 | <0.0001 |
| RS/SV+SV- vs SS/SV-SV- | 26.4 | 1.5-450.9 | 0.0237 |
| RR/SV+SV+ vs RS/SV+SV- | 15.0 | 6.9-32.6 | <0.0001 |
| **SV/CYP6P9a/CYP6P9b** |  |  |  |
| RR/RR/SV+SV+ vs SS/SS/SV-SV- | 374.7 | 21.6-6490.1 | <0.0001 |
| RS/RS/SV+SV- vs SS/SS/SV-SV- | 25.8 | 1.5-443.2 | 0.0248 |
| RR/RR/SV+SV+ vs RS/RS/SV+SV- | 15.2 | 6.9-33.5 | <0.0001 |

**S5 Table:**
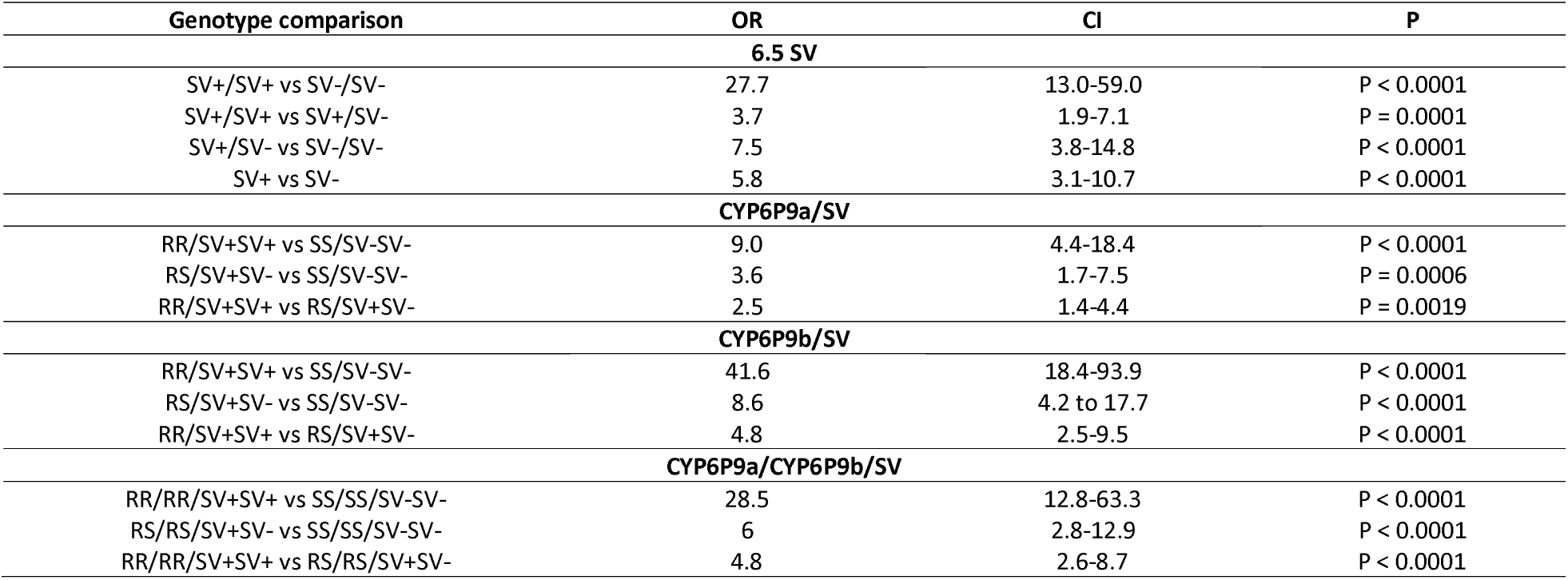
Correlation between genotypes of the 6.5kb SV and ability to survive exposure to PermaNet 2.0 in experimental huts using all samples.

| Genotype comparison | OR | CI | P |
| --- | --- | --- | --- |
| **6.5 SV** |  |  |  |
| SV+/SV+ vs SV-/SV- | 27.7 | 13.0-59.0 | P < 0.0001 |
| SV+/SV+ vs SV+/SV- | 3.7 | 1.9-7.1 | P = 0.0001 |
| SV+/SV- vs SV-/SV- | 7.5 | 3.8-14.8 | P < 0.0001 |
| SV+ vs SV- | 5.8 | 3.1-10.7 | P < 0.0001 |
| **CYP6P9a/SV** |  |  |  |
| RR/SV+SV+ vs SS/SV-SV- | 9.0 | 4.4-18.4 | P < 0.0001 |
| RS/SV+SV- vs SS/SV-SV- | 3.6 | 1.7-7.5 | P = 0.0006 |
| RR/SV+SV+ vs RS/SV+SV- | 2.5 | 1.4-4.4 | P = 0.0019 |
| **CYP6P9b/SV** |  |  |  |
| RR/SV+SV+ vs SS/SV-SV- | 41.6 | 18.4-93.9 | P < 0.0001 |
| RS/SV+SV- vs SS/SV-SV- | 8.6 | 4.2 to 17.7 | P < 0.0001 |
| RR/SV+SV+ vs RS/SV+SV- | 4.8 | 2.5-9.5 | P < 0.0001 |
| **CYP6P9a/CYP6P9b/SV** |  |  |  |
| RR/RR/SV+SV+ vs SS/SS/SV-SV- | 28.5 | 12.8-63.3 | P < 0.0001 |
| RS/RS/SV+SV- vs SS/SS/SV-SV- | 6 | 2.8-12.9 | P < 0.0001 |
| RR/RR/SV+SV+ vs RS/RS/SV+SV- | 4.8 | 2.6-8.7 | P < 0.0001 |

**S6 Table:** Correlation between genotypes of the 6.5kb SV and ability to survive exposure to PermaNet 3.0 in experimental huts using unfed samples.

| Genotype comparison | OR | CI | P |
| --- | --- | --- | --- |
| **6.5 SV** |  |  |  |
| SV+/SV+ vs SV-/SV- | 370.9 | 22.4-6151.4 | P < 0.0001 |
| SV+/SV+ vs SV+/SV- | 4.1 | 2.3-7.5 | P < 0.0001 |
| SV+/SV- vs SV-/SV- | 158.3 | 9.6-2619.9995 | P = 0.0004 |
| SV+ vs SV- | 3.8 | 1.9-7.6 | P = 0.0001 |
| **CYP6P9a/SV** |  |  |  |
| RR/SV+SV+ vs SS/SV-SV- | 423.6 | 25.5-7036.2 | P < 0.0001 |
| RS/SV+SV- vs SS/SV-SV- | 118.7 | 7.2-1967.5 | P = 0.0009 |
| RR/SV+SV+ vs RS/SV+SV- | 3.6 | 2.0-6.5 | P < 0.0001 |
| **CYP6P9b/SV** |  |  |  |
| RR/SV+SV+ vs SS/SV-SV- | 4.5 | 2.5-8.3 | P < 0.0001 |
| RS/SV+SV- vs SS/SV-SV- | 2.3 | 1.3-4.1 | P = 0.0062 |
| RR/SV+SV+ vs RS/SV+SV- | 2.0 | 1.1-3.6 | P = 0.0159 |
| **CYP6P9a/CYP6P9b/SV** |  |  |  |
| RR/RR/SV+SV+ vs SS/SS/SV-SV- | 423.6 | 25.5-7036.2 | P < 0.0001 |
| RS/RS/SV+SV- vs SS/SS/SV-SV- | 118.7 | 7.2-1967.5 | P = 0.0009 |
| RR/RR/SV+SV+ vs RS/RS/SV+SV- | 3.6 | 2.0-6.5 | P < 0.0001 |

**S7 Table:** Correlation between genotypes of the 6.5kb SV and ability to survive exposure to PermaNet 3.0 in experimental huts using all samples.

| Genotype comparison | OR | CI | P |
| --- | --- | --- | --- |
| **6.5 SV** |  |  |  |
| SV+/SV+ vs SV-/SV- | 443.5 | 26.7-7369.5 | P < 0.0001 |
| SV+/SV+ vs SV+/SV- | 4.1 | 2.3-7.5 | P < 0.0001 |
| SV+/SV- vs SV-/SV- | 108.9 | 6.6-1807 | P = 0.0011 |
| SV+ vs SV- | 3.8 | 1.9-7.6 | P = 0.0001 |
| **CYP6P9a/SV** |  |  |  |
| RR/SV+SV+ vs SS/SV-SV- | 511.3 | 30.7-8512.7 | P < 0.0001 |
| RS/SV+SV- vs SS/SV-SV- | 79.0 | 4.7-1315.5 | P = 0.0023 |
| RR/SV+SV+ vs RS/SV+SV- | 6.6122 | 3.6-12.3 | P < 0.0001 |
| **CYP6P9b/SV** |  |  |  |
| RR/SV+SV+ vs SS/SV-SV- | 511.3 | 30.7-8512.7 | P < 0.0001 |
| RS/SV+SV- vs SS/SV-SV- | 118.7 | 7.2-1967.5 | P = 0.0009 |
| RR/SV+SV+ vs RS/SV+SV- | 4.3 | 2.4-7.8 | P < 0.0001 |
| **SV/CYP6P9a/CYP6P9b** |  |  |  |
| RR/RR/SV+SV+ vs SS/SS/SV-SV- | 537.2 | 32.2-8950.6 | P < 0.0001 |
| RS/RS/SV+SV- vs SS/SS/SV-SV- | 82.9 | 5.0-1379.7 | P = 0.0021 |
| RR/RR/SV+SV+ vs RS/RS/SV+SV- | 6.6 | 3.6-12.7 | P < 0.0001 |

**S8 Table:** Correlation between genotypes of the 6.5kb SV and ability to blood-feed in the presence of PermaNet 2.0 in experimental huts

| Genotype comparison | OR | CI | P |
| --- | --- | --- | --- |
| **6.5 SV** |  |  |  |
| SV+/SV+ vs SV-/SV- | 2.1 | 1.2-3.8 | P = 0.0101 |
| SV+/SV+ vs SV+/SV- | 0.8 | 0.5-1.4 | P = 0.3956 |
| SV+/SV- vs SV-/SV- | 2.7 | 1.5-4.8 | P = 0.0007 |
| SV+ vs SV- | 1.3 | 0.8-2.3 | P = 0.3226 |
| **CYP6P9a/SV** |  |  |  |
| RR/SV+SV+ vs SS/SV-SV- | 4.1 | 2.3-7.4 | P < 0.0001 |
| RS/SV+SV- vs SS/SV-SV- | 2.7 | 1.5-4.8 | P = 0.0007 |
| RR/SV+SV+ vs RS/SV+SV- | 1.5 | 0.9-2.7 | P = 0.1480 |
| **CYP6P9b/SV** |  |  |  |
| RR/SV+SV+ vs SS/SV-SV- | 405 | 24.4-6723.3 | P < 0.0001 |
| RS/SV+SV- vs SS/SV-SV- | 265.7 | 16.1-4397.7 | P = 0.0001 |
| RR/SV+SV+ vs RS/SV+SV- | 1.5 | 0.9-2.7 | P = 0.1461 |
| **SV/CYP6P9a/CYP6P9b** |  |  |  |
| RR/RR/SV+SV+ vs SS/SS/SV-SV- | 2.3 | 1.3-4.0 | P = 0.0047 |
| RS/RS/SV+SV- vs SS/SS/SV-SV- | 1.5 | 0.9-2.7 | P = 0.1462 |
| RR/RR/SV+SV+ vs RS/RS/SV+SV- | 1.7 | 1.0-3.0 | P = 0.0620 |

## Notes

### Competing Interest Statement

The authors have declared no competing interest.

## References

1. Bhatt S, Weiss D, Cameron E, Bisanzio D, Mappin B, Dalrymple U, et al. The effect of malaria control on Plasmodium falciparum in Africa between 2000 and 2015. Nature. 2015;526(7572):207.

2. Barnes KG, Weedall GD, Ndula M, Irving H, Mzihalowa T, Hemingway J, et al. Genomic footprints of selective sweeps from metabolic resistance to pyrethroids in African malaria vectors are driven by scale up of insecticide-based vector control. PLoS Genet. 2017;13(2):e1006539.

3. WHO. World malaria report 2018. Geneva: WHO; 2018. 2018.

4. WHO. World Malaria Report 2019. Organization WH, editor 2019.

5. WHO. Global plan for insecticide resistance management in malaria vectors: executive summary. World Health Organization, 2012.

6. Martinez-Torres D, Chandre F, Williamson M, Darriet F, Berge JB, Devonshire AL, et al. Molecular characterization of pyrethroid knockdown resistance (kdr) in the major malaria vector Anopheles gambiae ss. Insect Mol Biol. 1998;7(2):179–84.

7. Irving H, Wondji CS. Investigating knockdown resistance (kdr) mechanism against pyrethroids/DDT in the malaria vector Anopheles funestus across Africa. BMC Genet. 2017;18(1):76.

8. Hemingway J, Ranson H. Insecticide resistance in insect vectors of human disease. Annu Rev Entomol. 2000;45(1):371–91.

9. Riveron JM, Yunta C, Ibrahim SS, Djouaka R, Irving H, Menze BD, et al. A single mutation in the GSTe2 gene allows tracking of metabolically-based insecticide resistance in a major malaria vector. Genome Biol. 2014;15(2):R27. doi: 10.1186/gb-2014-15-2-r27. PubMed PMID: 24565444.

10. Mitchell SN, Stevenson BJ, Müller P, Wilding CS, Egyir-Yawson A, Field SG, et al. Identification and validation of a gene causing cross-resistance between insecticide classes in Anopheles gambiae from Ghana. Proceedings of the National Academy of Sciences. 2012;109(16):6147–52.

11. Ibrahim SS, Riveron JM, Bibby J, Irving H, Yunta C, Paine MJ, et al. Allelic Variation of Cytochrome P450s Drives Resistance to Bednet Insecticides in a Major Malaria Vector. PLoS Genet. 2015;11(10):e1005618. doi: 10.1371/journal.pgen.1005618. PubMed PMID: 26517127; PubMed Central PMCID: PMCPMC4627800.

12. Lumjuan N, Rajatileka S, Changsom D, Wicheer J, Leelapat P, Prapanthadara L-a, et al. The role of the Aedes aegypti Epsilon glutathione transferases in conferring resistance to DDT and pyrethroid insecticides. Insect Biochem Mol Biol. 2011;41(3):203–9.

13. Weedall GD, Mugenzi LM, Menze BD, Tchouakui M, Ibrahim SS, Amvongo-Adjia N, et al. A cytochrome P450 allele confers pyrethroid resistance on a major African malaria vector, reducing insecticide-treated bednet efficacy. Science translational medicine. 2019;11(484):eaat7386.

14. Mugenzi LM, Menze BD, Tchouakui M, Wondji MJ, Irving H, Tchoupo M, et al. Cis-regulatory CYP6P9b P450 variants associated with loss of insecticide-treated bed net efficacy against Anopheles funestus. Nature communications. 2019;10(1):1–11.

15. Lucas ER, Miles A, Harding NJ, Clarkson CS, Lawniczak MK, Kwiatkowski DP, et al. Whole-genome sequencing reveals high complexity of copy number variation at insecticide resistance loci in malaria mosquitoes. Genome Res. 2019;29(8):1250–61.

16. Ingham VA, Pignatelli P, Moore JD, Wagstaff S, Ranson H. The transcription factor Maf-S regulates metabolic resistance to insecticides in the malaria vector Anopheles gambiae. BMC Genomics. 2017;18(1):669.

17. Wondji CS, Irving H, Morgan J, Lobo NF, Collins FH, Hunt RH, et al. Two duplicated P450 genes are associated with pyrethroid resistance in Anopheles funestus, a major malaria vector. Genome Res. 2009.

18. Amenya D, Naguran R, Lo TC, Ranson H, Spillings B, Wood O, et al. Over expression of a cytochrome P450 (CYP6P9) in a major African malaria vector, Anopheles funestus, resistant to pyrethroids. Insect Mol Biol. 2008;17(1):19–25.

19. Riveron JM, Irving H, Ndula M, Barnes KG, Ibrahim SS, Paine MJ, et al. Directionally selected cytochrome P450 alleles are driving the spread of pyrethroid resistance in the major malaria vector Anopheles funestus. Proceedings of the National Academy of Sciences. 2013;110(1):252–7.

20. Dickel DE, Visel A, Pennacchio LA. Functional anatomy of distant-acting mammalian enhancers. Philos Trans R Soc Lond B Biol Sci. 2013;368(1620):20120359. Epub 2013/05/08. doi: 10.1098/rstb.2012.0359. PubMed PMID: 23650633; PubMed Central PMCID: PMCPMC3682724.

21. Chung H, Bogwitz MR, McCart C, Andrianopoulos A, Batterham P, Daborn PJ. Cis-regulatory elements in the Accord retrotransposon result in tissue-specific expression of the Drosophila melanogaster insecticide resistance gene Cyp6g1. Genetics. 2007;175(3):1071–7.

22. Pennacchio LA, Bickmore W, Dean A, Nobrega MA, Bejerano G. Enhancers: five essential questions. Nat Rev Genet. 2013;14(4):288–95. Epub 2013/03/19. doi: 10.1038/nrg3458. PubMed PMID: 23503198; PubMed Central PMCID: PMCPMC4445073.

23. Riveron JM, Ibrahim SS, Mulamba C, Djouaka R, Irving H, Wondji MJ, et al. Genome-wide transcription and functional analyses reveal heterogeneous molecular mechanisms driving pyrethroids resistance in the major malaria vector Anopheles funestus across Africa. G3: Genes, Genomes, Genetics. 2017:g3. 117.040147.

24. Daborn P, Yen J, Bogwitz M, Le Goff G, Feil E, Jeffers S, et al. A single P450 allele associated with insecticide resistance in Drosophila. Science. 2002;297(5590):2253–6.

25. Catania F, Kauer M, Daborn P, Yen J, Ffrench-Constant R, Schlötterer C. World-wide survey of an Accord insertion and its association with DDT resistance in Drosophila melanogaster. Mol Ecol. 2004;13(8):2491–504.

26. Riveron JM, Huijben S, Tchapga W, Tchouakui M, Wondji MM, Tchoupo M, et al. Escalation of pyrethroid resistance in the malaria vector Anopheles funestus induces a loss of efficacy of PBO-based insecticide-treated nets in Mozambique. The Journal of infectious diseases. 2019. doi: 10.1093/infdis/jiz139. PubMed PMID: 30923819.

27. Riveron JM, Chiumia M, Menze BD, Barnes KG, Irving H, Ibrahim SS, et al. Rise of multiple insecticide resistance in Anopheles funestus in Malawi: a major concern for malaria vector control. Malar J. 2015;14(1):344. doi: 10.1186/s12936-015-0877-y. PubMed PMID: 26370361; PubMed Central PMCID: PMCPMC4570681.

28. Wilding CS, Smith I, Lynd A, Yawson AE, Weetman D, Paine MJ, et al. A cis-regulatory sequence driving metabolic insecticide resistance in mosquitoes: functional characterisation and signatures of selection. Insect Biochem Mol Biol. 2012;42(9):699–707. doi: 10.1016/j.ibmb.2012.06.003. PubMed PMID: 22732326.

29. Ibrahim SS, Ndula M, Riveron JM, Irving H, Wondji CS. The P450 CYP6Z1 confers carbamate/pyrethroid cross-resistance in a major African malaria vector beside a novel carbamate-insensitive N485I acetylcholinesterase-1 mutation. Mol Ecol. 2016;25(14):3436–52. doi: 10.1111/mec.13673. PubMed PMID: 27135886; PubMed Central PMCID: PMCPMC4950264.

30. Barnes KG, Irving H, Chiumia M, Mzilahowa T, Coleman M, Hemingway J, et al. Restriction to gene flow is associated with changes in the molecular basis of pyrethroid resistance in the malaria vector Anopheles funestus. Proc Natl Acad Sci U S A. 2017;114(2):286–91. doi: 10.1073/pnas.1615458114. PubMed PMID: 28003461; PubMed Central PMCID: PMCPMC5240677.

31. Barnes KG, Weedall GD, Ndula M, Irving H, Mzihalowa T, Hemingway J, et al. Genomic Footprints of Selective Sweeps from Metabolic Resistance to Pyrethroids in African Malaria Vectors Are Driven by Scale up of Insecticide-Based Vector Control. PLoS Genet. 2017;13(2):e1006539. doi: 10.1371/journal.pgen.1006539. PubMed PMID: 28151952; PubMed Central PMCID: PMCPMC5289422.

32. Michel AP, Ingrasci MJ, Schemerhorn BJ, Kern M, Le Goff G, Coetzee M, et al. Rangewide population genetic structure of the African malaria vector Anopheles funestus. Mol Ecol. 2005;14(14):4235–48. PubMed PMID: 16313589.

33. Wondji CS, Dabire RK, Tukur Z, Irving H, Djouaka R, Morgan JC. Identification and distribution of a GABA receptor mutation conferring dieldrin resistance in the malaria vector Anopheles funestus in Africa. Insect Biochem Mol Biol. 2011;41(7):484–91. Epub 2011/04/20. doi: S0965-1748(11)00080-4 [pii] 10.1016/j.ibmb.2011.03.012. PubMed PMID: 21501685.

34. Hunt R, Brooke B, Pillay C, Koekemoer L, Coetzee M. Laboratory selection for and characteristics of pyrethroid resistance in the malaria vector Anopheles funestus. Med Vet Entomol. 2005;19(3):271–5.

35. WHO. Test procedures for insecticide resistance monitoring in malaria vector mosquitoes. 2016.

36. WHO. Malaria entomology and vector control: World Health Organization; 2013.

37. Langmead B, Salzberg SL. Fast gapped-read alignment with Bowtie 2. Nat Methods. 2012;9(4):357–9. doi: 10.1038/nmeth.1923. PubMed PMID: 22388286; PubMed Central PMCID: PMCPMC3322381.

38. Wei Z, Wang W, Hu P, Lyon GJ, Hakonarson H. SNVer: a statistical tool for variant calling in analysis of pooled or individual next-generation sequencing data. Nucleic Acids Res. 2011;39(19):e132. doi: 10.1093/nar/gkr599. PubMed PMID: 21813454; PubMed Central PMCID: PMCPMC3201884.

39. Livak KJ. Organization and mapping of a sequence on the Drosophila melanogaster X and Y chromosomes that is transcribed during spermatogenesis. Genetics. 1984;107(4):611–34.

40. Hall TA, editor BioEdit: a user-friendly biological sequence alignment editor and analysis program for Windows 95/98/NT. Nucleic acids symposium series; 1999: [London]: Information Retrieval Ltd., c1979-c2000.

